# The brain detects stimulus features, but not stimulus conflict in task-irrelevant sensory input

**DOI:** 10.1101/596999

**Authors:** Stijn A. Nuiten, Andrés Canales-Johnson, Lola Beerendonk, Nutsa Nanuashvili, Johannes J. Fahrenfort, Tristan Bekinschtein, Simon van Gaal

## Abstract

Cognitive control over conflicting sensory input is central to adaptive human behavior. It might therefore not come as a surprise that past research has shown conflict detection in the absence of conscious awareness. This would suggest that the brain may detect conflict fully automatically, and that it can even occur without paying attention. Contrary to this intuition, we show that task-relevance is crucial for conflict detection. Univariate and multivariate analyses on electroencephalographic data from human participants revealed that when auditory stimuli are fully task-irrelevant, the brain disregards conflicting input entirely, whereas the same input elicits strong neural conflict signals when task-relevant. In sharp contrast, stimulus features were still processed, irrespective of task-relevance. These results show that stimulus properties are only integrated to allow conflict to be detected by prefrontal regions when sensory information is task-relevant and therefore suggests an attentional bottleneck at high levels of information analysis.

## Introduction

Every day we are bombarded with sensory information from the environment and we often face the challenge of selecting the relevant information and ignoring irrelevant –potentially conflicting– information to maximize performance. These selection processes require much effort and our full attention, sometimes rendering us deceptively oblivious to irrelevant sensory input (e.g. chest-banging apes), as illustrated by the famous inattentional blindness phenomenon (Simons & Chabris, 1999). However, unattended events that are not relevant for the current task might still capture our attention or interfere with ongoing task performance, for example when they are inherently relevant to us (e.g. our own name). This is for example illustrated in another famous psychological phenomenon: the cocktail party effect (Cherry, 1953; Moray, 1959). Thus, under specific circumstances, task-irrelevant information may capture attentional resources and be subsequently processed with different degrees of depth.

It is currently a matter of debate which cognitive processes require top-down attention and which not (Dehaene, Changeux, Naccache, Sackur, & Sergent, 2006; Koch & Tsuchiya, 2007; Koelewijn, Bronkhorst, & Theeuwes, 2010; Lamme, 2003; Rousselet, Thorpe, & Fabre-Thorpe, 2004; VanRullen, 2007). It was long thought that only basic physical stimulus features or very salient stimuli are processed in the absence of attention (Treisman & Gelade, 1980), due to an “attentional bottleneck” at higher-levels of analysis (Broadbent, 1958; Deutsch & Deutsch, 1963; Lachter, Forster, & Ruthruff, 2004; Wolfe & Horowitz, 2004). However, it has been shown recently that several tasks may in fact still unfold in the (near) absence of attention, including perceptual integration (Fahrenfort, van Leeuwen, Olivers, & Hogendoorn, 2017), the processing of emotional valence (Sand & Wiens, 2011; Stefanics, Csukly, Komlósi, Czobor, & Czigler, 2012), semantical processing of written words (Schnuerch, Kreitz, Gibbons, & Memmert, 2016) and visual scene categorization (Fei-Fei, VanRullen, Koch, & Perona, 2002; Gronau & Izoutcheev, 2017; Peelen, Fei-Fei, & Kastner, 2009). Although one should be cautious in claiming complete absence of attention (Lachter et al., 2004), these and other studies have pushed the boundaries of potential processing without attention, and may even question the existence of an attentional bottleneck at all. Perhaps, however, the attentional bottleneck is present, but at even higher levels of cognitive processing, i.e. cognitive control functions in prefrontal cortex. Here, we test whether such an attentional bottleneck indeed exists in the human brain and whether it is linked to neural signatures of prefrontal processing.

Specifically, we aim to test whether prefrontal cognitive control operations, necessary to identify and resolve conflict in sensory input, are operational when that input is fully irrelevant for the task at hand (and hence unattended). Previous work has shown that the brain has dedicated networks for the detection and resolution of conflict, in which the medial frontal cortex (MFC) plays a pivotal role (Ridderinkhof, Ullsperger, Crone, & Nieuwenhuis, 2004). Conflict detection and subsequent behavioral adaptation is central to human cognitive control and it is therefore maybe not surprising that past research has shown that conflict detection can even occur unconsciously (Atas, Desender, Gevers, & Cleeremans, 2016; D’Ostilio & Garraux, 2012a; Huber-Huber & Ansorge, 2018; van Gaal, Ridderinkhof, Fahrenfort, Scholte, & Lamme, 2008a), suggesting that the brain may detect conflict fully automatically and that it may even occur without paying attention (e.g. Rahnev, Huang, & Lau, 2012).

Here, we presented auditory spoken words (“left” and “right” in Dutch) through speakers located at the left and right side of the body. Crucially, by presenting these stimuli through either the left or the right speaker, content-location conflict arose on specific trials (e.g. the word “left” through the right speaker) but not on others (e.g. the word “right” through the right speaker). In the task-relevant condition, participants were instructed to respond according to the content of the stimulus, ignoring its location. In the task-irrelevant condition however, participants performed a visually demanding random dot-motion (RDM) task (discriminating upward/downward motion), while still being presented with the same auditory stimuli—now fully irrelevant for task performance. Behavioral responses on this visual task were orthogonal to the response tendencies potentially triggered by the auditory stimuli, excluding any task or response related interference (Fig. 1A). Electroencephalography (EEG) was recorded to test whether content and location information can be integrated to elicit conflict when stimuli are task-irrelevant. Specifically, we focused on conflict-induced theta-band neural oscillations over midfrontal electrodes, originating from the MFC (Cohen & Cavanagh, 2011; Cohen & Ridderinkhof, 2013; Cohen & van Gaal, 2013, 2014; Jiang, Zhang, & van Gaal, 2015; Nigbur, Cohen, Ridderinkhof, & Stürmer, 2012; Pastötter, Dreisbach, & Bäuml, 2013; Usami et al., 2013). These theta-band oscillations and behavioral conflict effects can undergo across-trial modulation, a phenomenon known as conflict adaptation (i.e. reduced conflict effects after conflict in the previous trial; Egner, 2007). We investigated whether these neural markers of conflict detection and adaptation could be observed for auditory conflicting stimuli when they were completely task-irrelevant. To maximize the possibility to observe conflict detection when conflicting stimuli are task-irrelevant, and to explore the effect of task automatization, participants performed the tasks both before and after extensive training, which may increase the efficiency of cognitive control (Fig. 1B) (van Gaal, Ridderinkhof, Fahrenfort, Scholte, & Lamme, 2008b).

**Figure 1.**
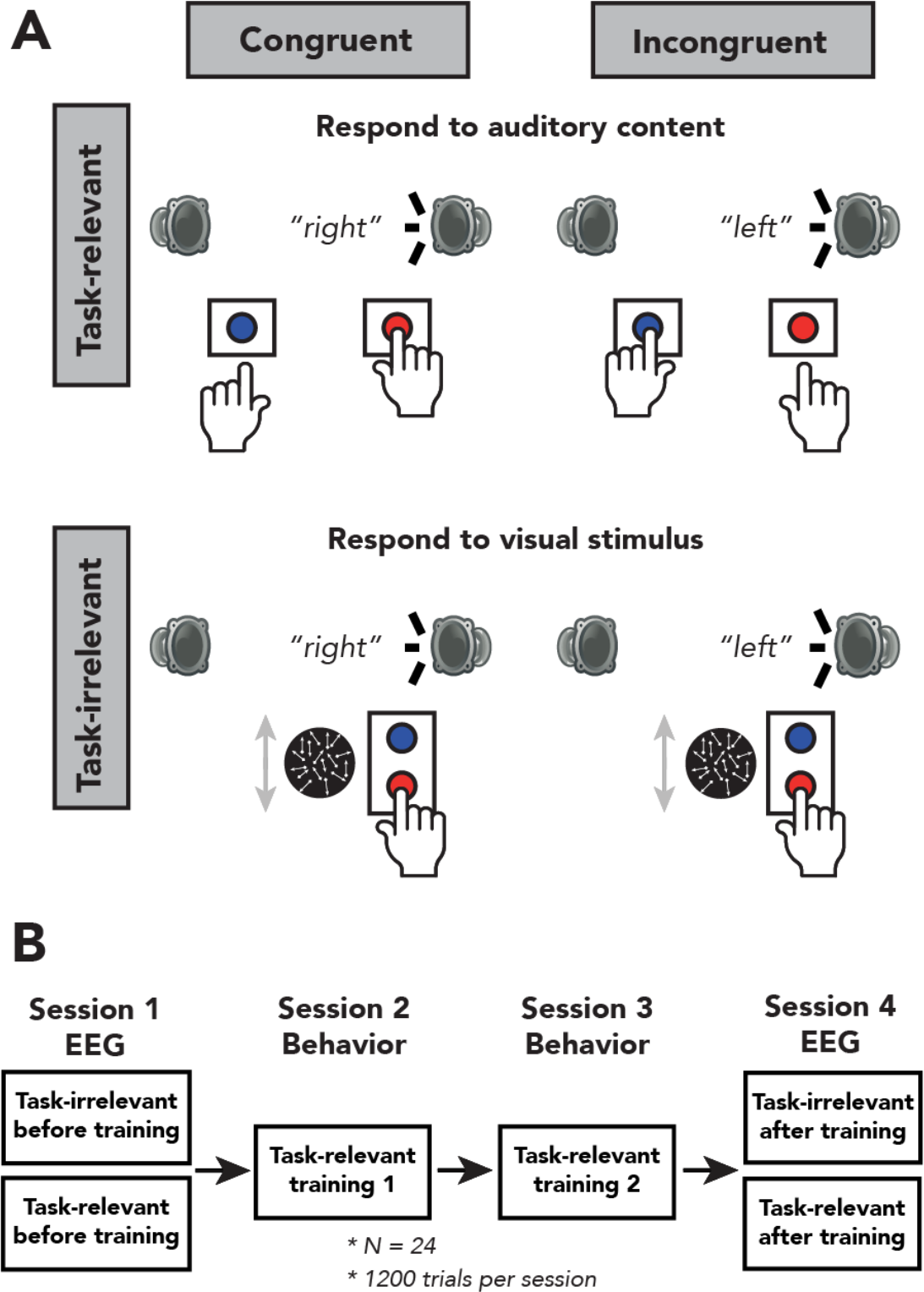
Experimental design. **(A)** Schematic representation of the experimental design for the task-relevant (top) and task-irrelevant (bottom) conditions. In both conditions, the spoken words “left” and “right were presented through either a speaker located on the left or right side of the participant. In this figure, sounds are only depicted as originating from the right, whereas in the experiment the sounds could also originate from the left speaker. In the task-relevant condition, participants were instructed to report the content (“left” or “right”) of an auditory stimulus via a button press with their left or right hand, respectively, and to ignore the spatial location at which the auditory stimulus was presented. During the task-irrelevant condition, participants were instructed to report the overall movement of dots (up or down) via a button press with their right hand, whilst still being presented with the auditory stimuli, which were therefore task-irrelevant. In both tasks, content of the auditory stimuli could be congruent or incongruent with its location of presentation (50% congruent/incongruent trials). **(B)** Overview of the sequence of the four experimental sessions of this study. Participants performed two EEG sessions during which they first performed the task-irrelevant condition followed by the task-relevant condition. Each session consisted of 1200 trials, divided over twelve blocks, allowing participants to rest in between blocks. In between experimental sessions, participants were trained on the task-relevant condition on two training sessions of 1hr each.

## Results

To investigate whether our experimental design was apt to induce conflict effects for task-relevant conflict and to test whether conflict effects were still present for task-irrelevant conflict, we performed repeated measures (rm-)ANOVAs (2×2×2×2 factorial) on mean reaction times (RTs) and error rates (ERs) gathered during the EEG-recording sessions (session 1, “before training”; session 4, “after training”). This allowed us to include 1) task-relevance (yes/no), 2) training (before/after), 3) congruency of auditory content with location of auditory source (congruent/incongruent) and 4) previous trial congruency as factors in our analyses. Behavioral data for all four sessions of the task-relevant condition are shown in figure 2B/D. Note that congruency is always defined based on the relationship between stimulus content (the word “left” or “right”) and location (left side, right side) of the auditorily presented stimuli, also when participants performed the visual task (and therefore the auditory stimuli were task-irrelevant).

**Figure 2.**
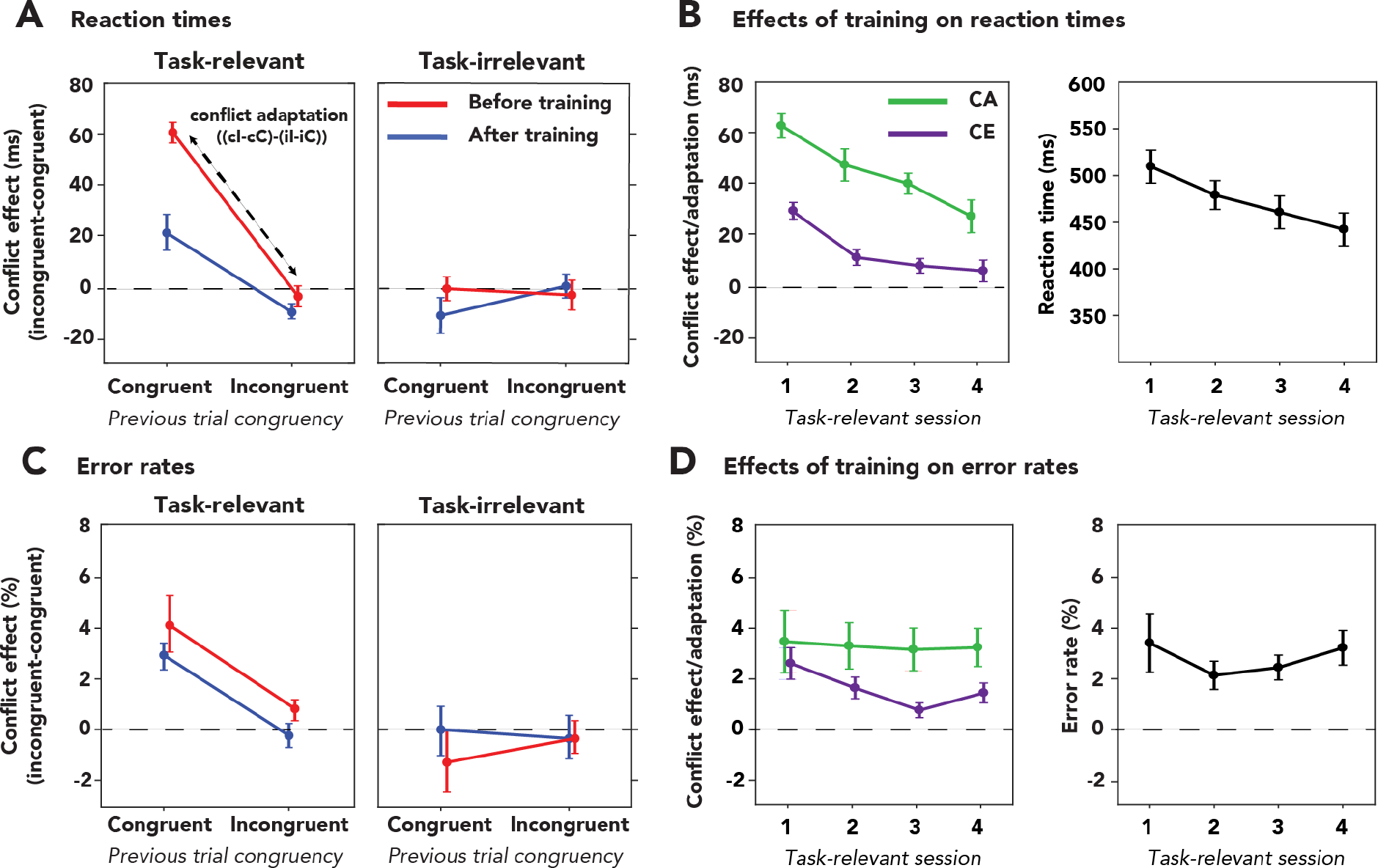
Behavioral results. **(A/C)** Current trial conflict effects (incongruent (uppercase I)– congruent (uppercase C)) and across-trial conflict adaptation effects –i.e. difference in the size of the conflict effect after a congruent previous trial (lowercase c) vs after a previous incongruent trial (lowercase i), reflected in downward slope in panel A and C (denoted as (cI-cC)-(iI-iC)) for the task-relevant (left) and task-irrelevant (right) conditions in reaction times (**A**) and error rates (**C**). Effects of conflict and conflict adaptation (difference in the size of the conflict effect after a congruent previous trial vs after a previous incongruent trial, reflected in downward slope in panel A and C) were only present when the auditory stimulus was task-relevant for both RTs and ERs. For the task-relevant condition conflict effects in terms of ERs and RTs were significantly smaller after training (blue line) as compared to before training (red line). Moreover, conflict adaptation effects also decreased following training, but only for RTs and not ERs. Error bars reflect the standard error of the mean. **(B/D)** Effect of training on conflict effect (CE, purple line) and conflict adaptation (CA, green line) in terms of RT (**B**) and ER (**D**) when conflict was task-relevant. In the right panels, grand average RT and ER are depicted. Although there is a clear training effect for RT (i.e. conflict effect and conflict adaptation), there is no effect of training on ER.

### Behavioral slowing and decreased accuracy only induced by task-relevant conflict

First we report the current trial conflict effects and thereafter we report across trial conflict adaption effects. Detection of conflict is typically associated with behavioral slowing and increased error rates. Indeed, we observed that participants were slower and made more errors on incongruent trials as compared to congruent trials (the conflict effect, RT: *F*_*1,23*_=8.58, *p*=0.008, 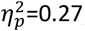; ER: *F*_*1,23*_=5.47, *p*=0.028, 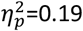). This conflict effect was modulated by task-relevance of the auditory stimuli (RT: *F*_*1,23*_=46.94, *p*<0.001, 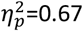; ER: *F*_*1,23*_=12.76, *p*=0.002, 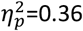) and post-hoc tests showed that the conflict effect was present when the auditory stimuli were task-relevant (RT_rel_: *F*_*1,23*_=38.56, *p*<0.001, 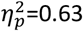; ER_rel_: *F*_*1,23*_=19.19, *p*<0.001, 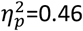), but not when they were task-irrelevant (RT_irrel_: *F*_*1,23*_=1.47, *p*=0.24, 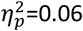, *BF*_*01*_=12.82; ER_irrel_: *F*_*1,23*_=1.00, *p*=0.33, 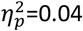, *BF*_*01*_=12.35, figure 2A/C). For the task-relevant condition the conflict effect decreased in magnitude after extensive training, suggesting increased efficiency of within-trial conflict resolution mechanisms (RT_rel_: *F*_*1,23*_=21.19, *p*<0.001, 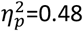; ER_rel_: *F*_*1,23*_=5.10, *p*=0.03, 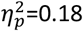, figure 2A/C). In general, in the task-relevant condition, mean RTs decreased after training, whereas ERs did not (RT_rel_: *F*_*1,23*_=35.51, *p*<0.001, 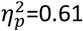; ER_rel_: *F*_*1,23*_=0.14, *p*=0.71, 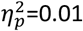, *BF*_*01*_=5.35, figure 2A/C). All other interactions of current trial effects were not reliable (*p*>0.05).

Conflict effects have been shown to undergo across-trial modulation (i.e. conflict adaptation) even when current or previous trial conflict is strongly masked (Jiang, Zhang, & van Gaal, 2015; van Gaal, Lamme, & Ridderinkhof, 2010). Because we did not observe current trial conflict effects of task-irrelevant conflict, no across-trial modulation was observed as well (RT_irrel_: *F*_*1,23*_=0.65, *p*=0.43, 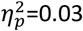, *BF*_*01*_=62.50; ER_rel_: *F*_*1,23*_=0.09, *p*=0.77, 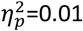, *BF*_*01*_=66.67). However, we did observe conflict adaptation for the task-relevant condition: the conflict effect was reduced when the previous trial was incongruent compared to congruent (RT_rel_: *F*_*1,23*_=133.96, *p*<0.001, 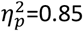; ER_rel_: *F*_*1,23*_=21.50, *p*<0.001, 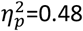; figure 2A/C). The overall magnitude of conflict adaptation was smaller after extensive training for RTs, but not for ERs (RT_rel_: *F*_*1,23*_=14.54, *p*<0.001, 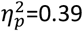; ER_rel_: *F*_*1,23*_=0.07, *p*=0.80, 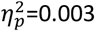, *BF*_*01*_=2.93, figure 2B/D). However, since the (current trial) conflict effect was strongly reduced after training (figure 2B), the relative magnitude (percentage) of conflict adaptation was in fact larger, showing increased efficiency of conflict adaptation as well (see supplementary figure S1 for more details).

### Midfrontal theta-band oscillations track task-relevant conflict detection and adaptation

The observation that task-irrelevant conflict has no effect on RTs and ERs, whereas task-relevant conflict does, is not surprising, because manual responses on the visual task (motion up/down with index and middle finger of right hand) were fully orthogonal to the conflicting nature of the auditory stimuli (left/right). Further, the task-irrelevant and task-relevant condition were tested in independent tasks, requiring different cognitive processes. For example, mean RTs on the task-irrelevant condition were on average 237ms longer than mean RTs for the task-relevant condition, see **supplementary information**). However, we should be cautious in concluding that there is no conflict detection of task-irrelevant input based on these behavioral results only, as neural effects can sometimes occur in absence of behavioral effects (e.g. Canales-Johnson et al., 2019 [preprint]; van Gaal et al., 2014). Therefore, in order to test whether unattended conflict is processed by the MFC conflict monitoring system we turn to neural data. We investigated whether specific neural markers of conflict processing, i.e. theta-band dynamics over medial frontal cortices, were present when conflict was task-relevant and/or task-irrelevant (Cohen & Donner, 2013; Cohen & van Gaal, 2013, 2014; Jiang, Zhang, & van Gaal, 2015; Nigbur et al., 2012; Wang, Li, Zheng, Wang, & Liu, 2014).

Replicating previous studies, current trial conflict (I-C, averaged over all conditions and two EEG-sessions) induced increased theta-band power at midfrontal electrodes (*p*<0.01, cluster-corrected; frequency range: 4Hz–8Hz, time range: 250ms–625ms). In line with several fMRI and animal studies, this conflict-related signal seemed to originate from the MFC (Botvinick, Cohen, & Carter, 2004; Ullsperger, Danielmeier, & Jocham, 2014; van Veen, Cohen, Botvinick, Stenger, & Carter, 2001, see source reconstructions in the inset of figure 3A). Band power was extracted from this time-frequency ROI in order to investigate effects of task-relevance, training, current trial and previous trial congruency using ANOVAs (figure 3B/D).

**Figure 3.**
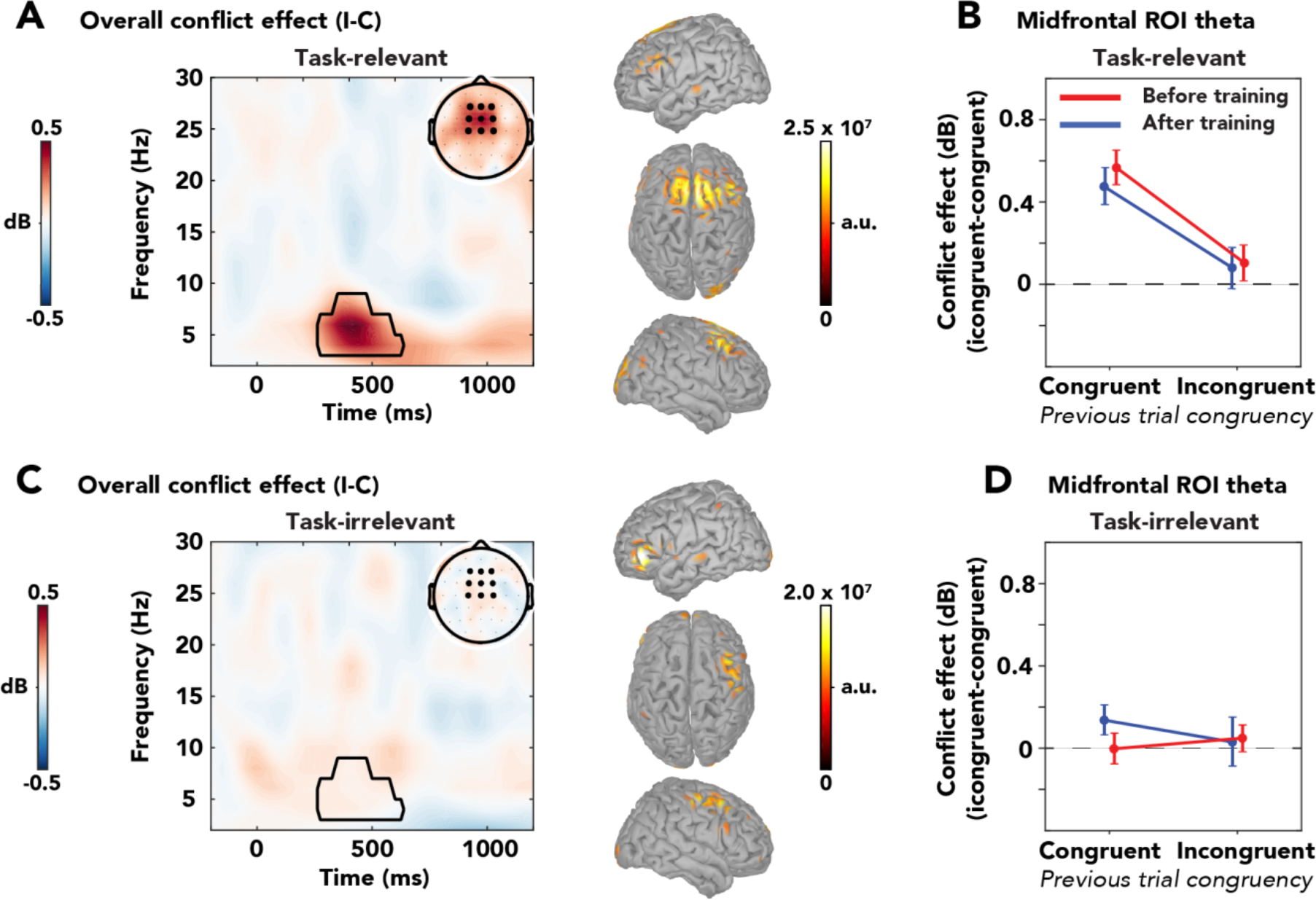
Conflict-induced midfrontal theta-band activity is present only when auditory stimuli are task-relevant. **(A/C)** Conflict effects in terms of time-frequency dynamics for the task-relevant **(A)** and task-irrelevant condition **(C)** over medial-frontal electrodes. The black delineated boxes indicate the time-frequency ROI in which overall conflict (I-C) was significant over conditions and sessions (corrected for multiple comparisons, **Methods**). Topographical distribution of oscillatory power within this time-frequency ROI is depicted in the top-right corner. Black dots represent the midfrontal EEG electrodes selected for obtaining the conflict-related theta-band power. A source-reconstruction analysis was performed on this time-frequency ROI; activations are expressed in arbitrary units (a.u.) and are thresholded at 50% of the maximum value of both conditions (Sergent, Baillet, & Dehaene, 2005; van Gaal et al., 2008b). Activations are depicted on unsmoothed brains, as reconstructed sources were only observed on the surface of the cortex. Note that the sources only serve a visualization purpose, as the method used to obtain them is sub-ideal and does not allow for proper statistical testing. **(B/D)** Conflict effects (I-C) and conflict adaptation effects ((cI-cC)-(iI-iC), reflected in downward slope) for the task-relevant **(B)** and task-irrelevant condition **(D)** of the conflict effect in dB (average ROI power I-C). Conflict and conflict adaptation effects were only present when the conflicting stimuli were task-relevant and were not affected by training.

The observed theta-band conflict effect was clearly modulated by task-relevance of the auditory stimuli as indicated by an interaction between current trial congruency and task-relevance (*F*_*1,23*_=23.42, *p*<0.001, 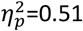, figure 3B/D). In fact, conflict-induced theta-band enhancements were only present when the auditory stimuli were task-relevant and not when they were task-irrelevant (theta_rel_: *F*_*1,23*_=35.23, *p*<0.001, 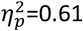; theta_irrel_: *F*_*1,23*_=1.72, *p*=0.20, 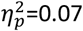). A Bayesian rm-ANOVA confirmed the absence of a theta-band conflict effect in the task-irrelevant condition (*BF*_*01*_=12.35). The observed theta-band conflict effect in the task-relevant condition was not affected by training of the conflicting task (*F*_*1,23*_=0.58, *p*=0.45, 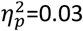, *BF*_*01*_=6.71).

Furthermore, theta-band conflict adaptation effects were only observed when the auditory stimulus was task-relevant (*F*_*1,23*_=28.73, *p*<0.001, 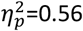) and were not modulated by training (*F*_*1,23*_=0.15, *p*=0.71, 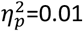, *BF*_*01*_=18.53). This indicates that, similar to behavior (RTs), theta-band conflict effects were smaller following incongruent trials versus congruent trials. All other main effects or interactions were not reliable (all *p*>0.05).

In sum, when the auditory stimuli were task-relevant and required a behavioral response that could be conflicting with the auditory content, effects of conflict detection and adaptation could be found in terms of reaction times, error rates and midfrontal theta-band oscillations. However, when these auditory stimuli were no longer task-relevant, and behavioral responses were not in conflict with the auditory material, these behavioral and neural markers of conflict vanished, even after considerable training on the task-relevant task. The latter conclusion was supported by Bayesian statistics providing support for the null hypothesis, indicating absence of conflict processing in the task-irrelevant condition.

### Multivariate confirmation: no detection of task-irrelevant conflict

Due to our hypothesis-driven analysis approach (selecting mid-frontal electrode channels) we may have missed neural dynamics modulated by conflict in the task-irrelevant task setting that did not occur in the hypothesized region. Plausibly, the neural dynamics of task-irrelevant conflict processing are spatially different from those related to the processing of task-relevant conflict. Therefore, in addition to the univariate analyses on the time-frequency data, we applied multivariate *decoding* techniques in the frequency domain to inspect in more detail whether and—if so—to what extent certain stimulus features were processed. These multivariate approaches have some advantages over traditional univariate approaches, for example, they are less sensitive to individual differences in spatial topography, because decoding accuracies are derived at a single subject level. Therefore, group statistics do not critically depend on the presence of effects in specific electrodes or clusters of electrodes.

We trained a classifier to distinguish between congruent vs. incongruent trials, for both the task-relevant and task-irrelevant condition. Above-chance classification accuracies imply that relevant information about the decoded stimulus feature is present in the neural data, meaning that some processing of that feature occurred (Hebart & Baker, 2017). We first performed our analysis on data from both EEG sessions separately to investigate effects of training. However, decoding accuracies did not differ significantly between sessions (p>0.05, cluster-corrected; see supplementary figure S2). Therefore, we subsequently conducted the decoding analysis on merged data from before and after the extensive behavioral training in the conflict task, thereby maximizing power to establish effects in our crucial comparisons. Thus, results reported here were obtained through the multivariate analyses of EEG-data of the two sessions combined.

Decoding of congruency yielded similar results as the univariate approach; stimulus congruency was represented in neural data only when conflict was task-relevant (*p*<0.05, cluster-corrected; frequency-range: 2-14Hz, peak frequency: 6Hz, time-range: 188-672ms, peak time: 375ms; figure 4A). Similar to our univariate results, conflict was mainly processed by the MFC, which further confirmed our *a priori* defined spatial region of interest (electrode selection) in the univariate analysis (see source reconstructions in the inset of figure 4A). Assessment of the qualitative difference in decoding performance between the task-relevant and task-irrelevant conditions resulted in two significant clusters of increased classifier accuracy when conflict was task-relevant: a theta-band cluster (frequency-range: 2-8Hz, peak frequency: 6Hz, time range: 250-672ms, peak time: 344ms, *p*=0.001 cluster-corrected; figure 4A) and an alpha-band cluster (frequency-range: 10-14Hz, peak frequency: 12Hz, time range: 469-672ms, peak time 547ms, p=0.017 cluster-corrected; figure 4A). A Bayesian one-sample t-test (one-sided, 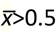) provided (moderate) evidence in favor of the null-hypothesis that task-irrelevant conflict is not processed by the brain (*BF*_*01*_=9.13, see **Methods**).

**Figure 4.**
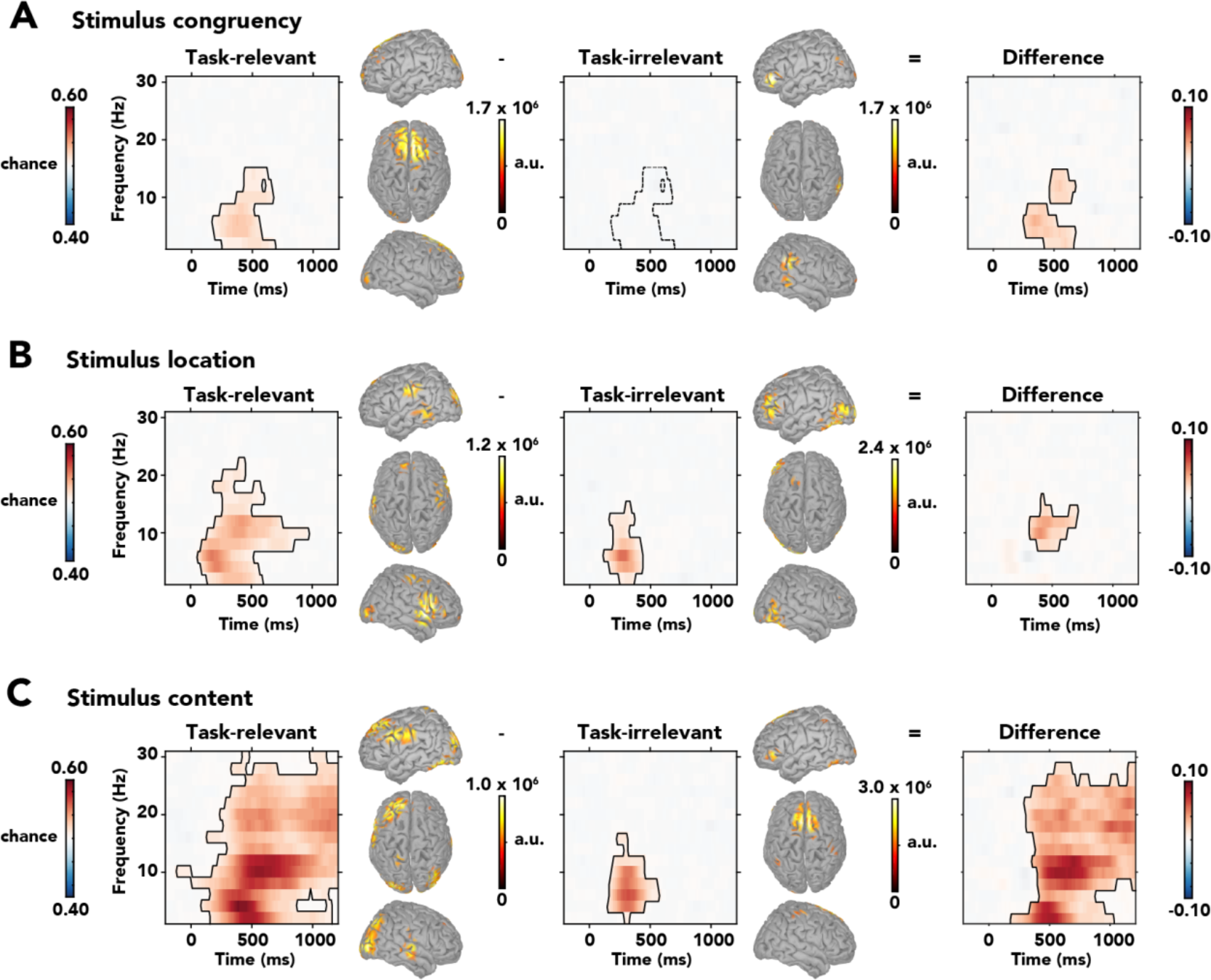
Classifier accuracies for congruency, location and content. The accuracies are depicted across the frequency-range (2-30Hz) for the task-relevant condition (left panels), task-irrelevant condition (middle panels) and their difference (right panels; relevant-irrelevant). Classifier accuracy is thresholded (cluster-base corrected, p<0.05) and significant clusters are outlined with a solid black line. Source activations are expressed in arbitrary units (a.u.) and are thresholded at 50% of the within-condition maximum value. **(A)** Classifier accuracies for stimulus congruency. Information about congruency was only present in the task-relevant condition. The dashed line in the middle panel reflects the mask from the task-relevant condition that was used to extract classifier weights for the sources of the task-irrelevant condition in panel **A**. **(B)** Classifier accuracies for stimulus location. Location of the auditory stimulus could be decoded in both conditions, meaning that information about this stimulus feature is present in neural signals. In the task-relevant condition, there was significantly more information about stimulus location represented in an alpha-band cluster. **(C)** Classifier accuracies for stimulus content. Content of the auditory stimulus could be decoded, and were thus processed by the brain, in both conditions. Information about stimulus content was however represented more long-lasting and across the spectral range when the auditory stimulus was task-relevant vs. when it was task-irrelevant.

### Stimulus features are processed in parallel, independent of task-relevance

Thus, prefrontal control networks no longer appear to detect conflict when it is task-irrelevant. However, is this observation specific to the conflicting nature of the auditory stimuli, or were the auditory stimuli not processed whatsoever when attention was reallocated in the visually demanding task? To address this question, we trained classifiers on two features of the auditory stimuli, i.e. location and content, to test whether these features are processed by the brain regardless of task-relevance. Indeed, the location of auditory stimuli was processed both when the stimuli were task-relevant (*p*<0.05, cluster-corrected; frequency-range: 2-22Hz, peak frequency: 6Hz, time-range: 63-953ms, peak time: 188ms; figure 4B) and task-irrelevant (*p*<0.05, cluster-corrected; frequency-range: 2-14Hz, peak frequency: 6Hz, time-range: 125-422ms, peak time: 281ms; figure 4B). The reconstructed sources of these effects were distributed across the cortex, including more frontal areas (see source reconstructions in the inset of figure 4B). When the auditory stimuli were task-relevant, however, increased classifier performance was present in an alpha-band cluster, as compared to when the stimuli were task-irrelevant (frequency-range: 8-16Hz, peak frequency: 10Hz, time-range: 313-703ms, peak time: 406ms, *p*=0.001 cluster-corrected; figure 4B).

Similarly, the content of the auditory stimuli could also be decoded from neural data for both the task-relevant (*p*<0.05, cluster-corrected; frequency-range: 2-30Hz, peak frequency: 4Hz, time-range: −125-1203ms, peak time: 391ms; figure 4C) and task-irrelevant condition (*p*<0.05, cluster-corrected; frequency-range: 2-16Hz, peak frequency: 6Hz, time-range: 156-578ms, peak time: 328ms; figure 4C). The content of task-relevant stimuli was processed by a globally distributed network, whereas results were more spatially confined for task-irrelevant stimuli (see source reconstructions in the inset of figure 4C). For the task-relevant condition, stimulus content was represented in more sustained and broadband neural activity as compared to the task-irrelevant condition (*p*<0.001, cluster-corrected; frequency-range: 2-28Hz, peak frequency: 10Hz, time-range: 188-1203ms, peak time: 578; figure 4C). The above chance performance of the classifiers for the stimulus features of the auditory stimulus demonstrates that location and content were processed, even when the auditory stimuli were task-irrelevant. Processing of stimulus features of task-irrelevant stimuli was, however, more transient in time, less distributed spatially, and more narrow-band in frequency as compared to processing of the same features of task-relevant stimuli.

Moreover, the content of the auditory stimulus could also be decoded from data obtained in the first session of the task-irrelevant condition, when participants had not yet performed the conflicting task (see supplementary figure S4). During this first session, the auditory stimuli had therefore not been coupled to any response-mappings whatsoever, thereby excluding potential effects of acquired task-relevance of these stimuli as a result of frequent exposure to the conflict task. Thus, task-related response-mappings belonging to the conflict task do not explain our observations of feature extraction in the task-irrelevant condition. Summarizing, we show that when conflicting stimuli are truly and consistently task-irrelevant, they are probably no longer detected by –nor relevant to– the MFC conflict monitoring system, but their features (content and location) are still extracted.

In conclusion, we report processing of stimulus features (i.e. location and content of auditory stimulus) regardless of task-relevance of the auditory stimulus, but a lack of integration of these features to form conflict when the auditory stimulus was task-irrelevant.

## Discussion

Although it has been hypothesized for a long time that only basic physical properties of task-irrelevant sensory input are processed (Treisman & Gelade, 1980), over the past few years a plethora of processes have been found to still occur in the absence of attention (e.g. Fahrenfort et al., 2017; Fei-Fei et al., 2002; Sand & Wiens, 2011). Here we aimed to push the limits and address the question as to whether prefrontal cognitive control networks could be recruited when conflicting sensory input is task-irrelevant. Interestingly, similar cognitive control functions have been shown to occur when stimuli are masked to be rendered unconscious (Atas et al., 2016; D’Ostilio & Garraux, 2012; Huber-Huber & Ansorge, 2018; van Gaal et al., 2008).

Participants performed an auditory-spatial conflict task (Buzzell, Roberts, Baldwin, & McDonald, 2013), in which conflicting auditory stimuli (e.g. the word “left” presented on the right side) were task-relevant in one task and task-irrelevant in another. When the auditory stimuli were task-relevant, we observed clear signals of conflict processing in behavior (i.e. longer reactions times and increased error rates) and brain activity (i.e. enhanced midfrontal theta-band power), replicating previous studies (Cavanagh & Frank, 2014; Cohen & Ridderinkhof, 2013; Jiang, Zhang, & van Gaal, 2015; Pastötter et al., 2013). Decoding of time-frequency data confirmed the modulation of midfrontal theta-band oscillations during conflict processing, as congruency could be classified within this frequency band, centered over the MFC. On the contrary, when the auditory stimuli were task-irrelevant all signs of conflict detection, in behavior and brain activity, vanished. Strikingly, the stimulus features, i.e. stimulus location/content, that ultimately need to be combined to generate conflict, were still processed, regardless of task-relevance. These results highlight that relatively basic stimulus properties escape the attentional bottleneck, lending support to previous studies (e.g. Fahrenfort et al., 2017; Fei-Fei et al., 2002; Sand & Wiens, 2011; Treisman & Gelade, 1980), but furthermore showcase that the bottleneck exists higher up the hierarchy of cognitive processing. We will interpret these results in more detail below.

### Task-irrelevant stimuli versus task-irrelevant stimulus features

Contrary to the current study, previous studies using a variety of conflict-inducing paradigms and attentional manipulations did report conflict effects in behavior and electrophysiological recordings induced by unattended stimuli or stimulus features (Mao & Wang, 2008; Padrão, Rodriguez-Herreros, Pérez Zapata, & Rodriguez-Fornells, 2015; Zimmer, Itthipanyanan, Grent-’T-Jong, & Woldorff, 2010). However, our study deviates from those studies in several crucial aspects. First, behavioral responses to the visual stimuli (response: up/down with index and middle finger of right hand) in our task-irrelevant condition were fully orthogonal to the stimulus features of the auditory stimuli (left/right), rendering the auditory stimuli truly task-irrelevant. In previous studies however, at least one stimulus feature was always associated with a response, thus the stimuli were not truly irrelevant for task performance (although some stimulus features were), therefore the stimuli were attended in some dimension of relevance. For example, Zimmer et al. (2010) have reported multisensory conflict effects in EEG, induced by to-be-ignored spoken letters that were congruent or incongruent with visually presented, task-relevant, letters (e.g. the letter ‘h’ presented both visually and auditorily: no conflict). The overlap in response mappings between auditorily and visually presented letters, however, rendered both stimuli task-relevant to some extent. One might argue that the same holds for our task-irrelevant condition, as the auditorily presented stimuli could potentially conflict with responses to the RDM-task that were exclusively made with the right hand, e.g. the spoken word “left” or the sound from left location may conflict generally more with a right hand response (independent of the up/down classification) than the spoken word “right” or the sound from right location. However, this was not the case, as both stimulus content and location did not affect mean RTs in the task-irrelevant condition, confirming that the auditorily presented stimuli were truly task-irrelevant (see supplementary figure S3 for details).

Second, in other studies (conflicting) stimuli were often task-irrelevant on one trial (e.g. because it is presented at an unattended location) but task-relevant on the next (e.g. because they were presented at the attended location) (e.g. Padrão et al., 2015; Zimmer et al., 2010). Such trial-by-trial fluctuations of task-relevance allow for across-trial modulations to confound any current trial effects (e.g. conflict-adaptation effect) and also induce some sort of *stand-by* attentional mode where participants never truly disengage to be able to determine if a stimulus is task-relevant. We prevented such confounding effects, as in the present study conflict information was task-irrelevant *on every single trial* in the task-irrelevant condition.

### Difference between response and perceptual conflict cannot account for absence of conflict detection

As the auditory stimuli were fully task-irrelevant in our paradigm, the conflict was perceptual, or intrinsic, in nature, opposed to the response conflict induced in previous studies (Kornblum, 1994). Although perceptual conflict effects are usually weaker than response conflict effects, both in behavior and electrophysiology (Frühholz, Godde, Finke, & Herrmann, 2011; van Veen et al., 2001; Wang et al., 2014), this difference in the origin of the conflict is unlikely to explain why we did not observe effects of task-irrelevant conflict, as opposed to earlier studies. First, several neurophysiological studies have previously reported electrophysiological modulations by perceptual conflict over the MFC (Jiang, Zhang, & Van Gaal, 2015; Nigbur et al., 2012; van Veen et al., 2001; Wang et al., 2014; Zhao et al., 2015). Second, an earlier study using a task very similar to ours (but only task-relevant stimuli) showed effects of perceptual conflict, i.e. unrelated to response codes, in terms of behavioral and neural data (Buzzell et al., 2013). Third, the prefrontal monitoring system has previously been observed to respond when subjects view other people making errors (Jääskeläinen et al., 2016; Van Schie, Mars, Coles, & Bekkering, 2004), suggesting that cognitive control can be triggered without the need to make a response. Similarly, during the task-irrelevant condition, the perceptually conflicting auditory stimuli (e.g. “left” coming from right location) could theoretically have triggered cognitive control, but the absence of conflict-related effects suggests that task-relevance is indeed necessary for prefrontal regions to detect such conflict.

### Inattentional deafness or genuine processing of stimulus features?

Besides the hypothesized conflict-related medial frontal theta-oscillations, we also report neural activity related to the processing of stimulus location and content. The lack of conflict effects in behavior and EEG when auditory stimuli were task-irrelevant might, at first glance, suggest a case of inattentional deafness, a phenomenon known to be induced by demanding visual tasks, which manifests itself in weakened early (~100ms) auditory evoked responses (Molloy, Griffiths, Chait, & Lavie, 2015). Interestingly, human speech seems to escape such load modulations and is still processed when unattended and task-irrelevant, potentially because of its inherent relevance, similar to (other) evolutionary relevant stimuli such as faces (Finkbeiner & Palermo, 2017; Lavie, Ro, & Russell, 2003; Röer, Körner, Buchner, & Bell, 2017; Zäske, Perlich, & Schweinberger, 2016). Indeed, the results of our multivariate analyses demonstrate that spoken words are processed (at least to some extent) when they are task-irrelevant, as stimulus content (the words “left” and “right”) and stimulus location (whether the word was presented on the left or the right side) could be decoded from time-frequency data (figure 4B/C, middle panels). More specifically, in terms of decoding accuracy, early processing (<300ms) of stimulus content and location was virtually identical for task-relevant and task-irrelevant auditory stimuli (figure 4C, right panel). For early processing of stimulus content, classification was most accurate in the theta-band, with peak frequencies of 4Hz (task-relevant) and 8Hz (task-irrelevant), which is in line with a recently proposed theoretical role for theta-oscillations in speech processing, namely that they track the acoustic-envelope of speech (Giraud & Poeppel, 2012). After this initial processing stage, processing of stimulus content is reflected in more durable and broadband activity for task-relevant auditory stimuli as compared to task-irrelevant stimuli, possibly related to higher-order processes (e.g. conflict resolution) and motor response preparation (figure 4C).

Early processing of stimulus location was also most strongly reflected in the theta-band (peaking at ~6Hz, figure 4B), which may relate to the auditory N1 ERP component, an ERP signal that is modulated by stimulus location (Fuentemilla, Marco-Pallarés, & Grau, 2006; Lewald & Getzmann, 2011; Salminen et al., 2015). Moreover, for task-relevant auditory stimuli, processing of stimulus location was significantly stronger in the alpha-band around 300-700ms post-stimulus (figure 4B, right panel), convergent with the previously reported role of alpha-band oscillations in orienting and allocating audio-spatial attention (Weisz, Müller, Jatzev, & Bertrand, 2014).

Thus, the characteristics of the early sensory processing of (task-irrelevant) auditory stimulus features are in line with recent findings of auditory and speech processing. Moreover, our observations correspond with recent empirical findings that suggest a dominant role for late control operations, as opposed to early selection processes, in resolving conflict (Itthipuripat, Deering, & Serences, 2019). Specifically, this work showed that in a Stroop-like paradigm both target and distractor information is analyzed fully, after which prefrontal regions resolve the conflicting input. Extending on this, we show intact initial processing of task-irrelevant sensory input, but deteriorated late control operations necessary to detect conflict, at least in the current set-up.

### Task-relevance as a prerequisite for conflict detection

So why are stimulus content and location not integrated into conflict by the MFC when the auditory stimulus is task-irrelevant? It has been proposed that the MFC monitors conflict through the detection of coinciding inputs (for theoretical model see: Cohen, 2014). In our paradigm, these afferents would carry information about the content and location, which have to be processed to some extent before this information can be transmitted to the MFC conflict monitoring system. This process may consist of multiple processing steps, possibly requiring different cortical areas to communicate with each other. Speculatively, the relatively fleeting temporal characteristics of the observed processing of the task-irrelevant stimulus features might have prevented their integration simply due to a lack of time. However, the time-window in which conflict was decodable in the task-relevant condition (188-672ms) coincides with the time-range in which stimulus location (125-422ms) and stimulus content (156-578ms) could be decoded during the task-irrelevant condition, suggesting that integration of these features could have occurred anywhere in this time-window. Therefore, it seems unlikely that the more temporally constrained processing of task-irrelevant stimulus features is the cause of deteriorated conflict integration. Alternatively, processing of these features may have been constrained to early sensory cortices and simply did not progress to frontal cortices, including the MFC, necessary for the detection of conflict. However, our source-reconstruction analysis of the multivariate data seems to dismiss this notion since more anteriorly located cortical areas seemed to be involved in the processing of the stimulus features, even in the absence of task-relevance. Importantly, previous work of our group demonstrated that masked task-irrelevant conflicting cues induce similar early processing in sensory cortices as compared to masked task-relevant cues, but prohibit activation of frontal cortices (van Gaal et al., 2008b). These seemingly opposing findings highlight that uncovering the role of task-relevance in processing of unconscious information deserves more attention in future work (see also van Gaal, de Lange, and Cohen, 2012).

### Automization of conflict processing does not promote detection of task-irrelevant conflict

Previous studies investigating unconscious conflict through masking procedures concluded that conflict detection by the MFC is automatic and does not require conscious awareness of the stimulus (D’Ostilio & Garraux, 2012; Jiang et al., 2015; van Gaal et al., 2008). Such automaticity can often be enhanced through training of the task. For example, after training of a semantic categorization lexical decision making task, response preparation signals were detected over the motor cortex of sleeping participants when new task-related words were presented (Kouider, Andrillon, Barbosa, Goupil, & Bekinschtein, 2014). Moreover, training in a stop-signal paradigm in which stop-signals were rendered unconscious through masking, led to an increase in the effectiveness of these stimuli to affect behavior (van Gaal, Ridderinkhof, van den Wildenberg, & Lamme, 2009). In order to see whether this automaticity could be enhanced in our paradigm, thereby hypothetically increasing the likelihood of conflict detection, we included training sessions of the conflict task in between EEG-recording sessions. However, even after this extensive training procedure (i.e. 3600 trials) no signs of conflict detection, in terms of behavior and mid-frontal theta dynamics, were present when the auditory stimuli were task-irrelevant. Training did result in decreased conflict effects in the task-relevant condition, indicating that our training procedure was successful and participants improved both within-trial conflict resolution mechanisms as well as across-trial conflict adaptation (see supplementary figure S1), suggesting more efficient conflict resolution mechanisms in general. Therefore, we can conclude that the automaticity of conflict detection by the MFC does not hold when auditory conflict is task-irrelevant (at least after the extent of training as studied here: three sessions).

### Conclusion

Summarizing, high-level cognitive processing that requires the integration of lower-level stimulus features seems to be diminished when stimuli are fully task-irrelevant, contrary to previous findings of perceptual processing outside the scope of attention (Peelen et al., 2009; Sand & Wiens, 2011; Schnuerch et al., 2016; Tusche, Kahnt, Wisniewski, & Haynes, 2013). These findings suggest crucial limitations of the brain’s capacity to process task-irrelevant “complex” cognitive control-initiating stimuli, suggestive of an attentional bottleneck at high levels of information analysis. In contrast, in absence of task-relevance, the processing of more basic physical features of sensory input appears to be less severely deteriorated (Lachter et al., 2004).

## Methods and Materials

### Participants

24 healthy human participants (18 female) aged 18 to 30 (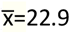, s=3.2), recruited from the University of Amsterdam (Amsterdam, The Netherlands), participated in this experiment for monetary compensation or participant credits. All participants had normal or corrected-to-normal vision and had no history of head injury or physical and mental illness. This study was approved by the local ethics committee of the University of Amsterdam and written informed consent was obtained from all participants after explanation of the experimental protocol.

### Design and procedures

Participants performed two tasks in which conflicting auditory stimuli were either task-relevant or task-irrelevant. In both tasks, conflict was elicited through a paradigm adapted from Buzzell et al. (2013), in which spatial information and content of auditory stimuli could interfere. In the task-relevant condition participants had to respond to the auditory stimuli, whereas in the task-irrelevant condition participants had to perform a demanding RDM-task, while the auditory conflicting stimuli were still presented (figure 1A). Participants performed both tasks on two experimental sessions of approximately 2.5 hours. In between these two experimental sessions, participants had two training sessions of one hour during which they only performed the task-relevant task (figure 1B). On each experimental session, participants first performed a shortened version of the RDM-task to determine the appropriate coherency of the moving dots (73%-77% correct), followed by the actual task-irrelevant condition and finally the task-relevant condition. Participants were seated in a darkened, sound isolated room, 50cm from a 69×39cm screen (frequency: 120Hz, resolution: 1920 × 1080, RGB: 128, 128, 128). Both experiments were programmed in MATLAB (R2012b, The MathWorks, Inc.), using functionalities from Psychtoolbox (Kleiner, Brainard, & Pelli, 2007).

### Task-relevant conflict condition

In the task-relevant condition, the spoken words “links” (i.e. “left” in Dutch) and “rechts” (i.e. “right” in Dutch) were presented through speakers located on both sides of the participant (figure 1A). Auditory stimuli were matched in duration and sample rate (44 kHz) and were recorded by the same male voice. By presenting these stimuli through either the left or the right speaker, content-location conflict arose on specific trials (e.g. the word “left” through the right speaker). Trials were classified accordingly, as either congruent (i.e. location and content are the same) or incongruent (i.e. location and content are different). Participants were instructed to respond as fast and accurate as possible, by pressing left (“a”) or right (“l”) on a keyboard located in front of the participants, according to the stimulus content, ignoring stimulus location. Responses had to be made with the left or right index finger, respectively. The task was divided in twelve blocks of 100 trials each, allowing participants to rest in between blocks. After stimulus presentation, participants had a 2s period in which they could respond. A variable inter-trial-interval between 850ms-1250ms was initiated directly after the response. If no response was made, the subsequent trial would start after the 2s response period. Congruent and incongruent trials occurred equally often (i.e. 50% of all trials), as expectancy of conflict has been shown to affect conflict processing (Soutschek, Stelzel, Paschke, Walter, & Schubert, 2015). Due to an error in the script, there was a disbalance in the amount of trials coming from the left (70%) vs. right (30%) speaker location for the first 14 participants. However, the amount of congruent vs. incongruent and “left” vs. “right” trials were equally distributed. For the upcoming analyses, all trial classes were balanced in trial count.

### Task-irrelevant conflict condition

In the condition where auditory stimuli were task-irrelevant, participants performed a RDM-task in which they had to discriminate the motion (up versus down) of white dots (n=603) presented on a black circle (RGB: 0, 0, 0; ~14° visual angle; see figure 1A). Onset of the visual stimulus was paired with the presentation of the auditory conflicting stimulus. Participants were instructed to respond according to the direction of the dots, by pressing the “up” or “down” key on a keyboard with their right hand as fast and accurate as possible. Again, participants could respond in a 2s time-interval which was terminated after responses and followed by an inter-trial-interval of 850-1250ms. Task difficulty, in terms of dot-motion coherency (i.e. proportion of dots moving in the same direction) was titrated between blocks to 73-77% correct of all trials within that block. Similar to the task-relevant condition, the task-irrelevant condition was divided in twelve blocks containing 100 trials each, separated by short breaks. Again, congruent and incongruent trials, with respect to the auditory stimuli, occurred equally often.

### Data analysis

We were primarily interested in the effects of congruency of the auditory stimuli on both behavioral and neural data. Therefore, we defined trial-congruency on the basis of these auditory stimuli, both in the task-relevant and the task-irrelevant condition. Thus, the direction of dot-motion in the task-irrelevant condition was disregarded in the data-analysis.

### Behavioral data analysis

The first trial of every block, incorrect or missed trials, trials following incorrect responses and trials with an RT <100ms or >1500ms were excluded from behavioral analyses (17.7% of all trials). Conflict on trial *n* has been found to cause increased ERs and prolonged RTs, as compared to trials with no conflict. This current trial effect of conflict can be modulated by previously experienced conflict on trial *n-1*, a phenomenon called conflict adaptation. In order to investigate whether current trial conflict effects and the modulation thereof by previous conflict were present both when conflict was task-relevant and task-irrelevant and to inspect if training of the conflict task affected such conflict effects, we performed rm-ANOVAs on ERs and RTs with task-relevance, training (before vs. after), current trial congruency and previous trial congruency as factors (2×2×2×2 factorial design). Additional post-hoc rm-ANOVAs, for the task-relevant and task-irrelevant conditions separately (2×2×2 factorial design), were used to inspect the origin of significant factors that were modulated by task-relevance. In case of null findings, we performed a Bayesian rm-ANOVA with similar factors, to verify if there is actual support of the null hypothesis (JASP Team, 2018).

### EEG: recording and preprocessing

EEG-data were recorded with a 64-channel BioSemi apparatus (BioSemi B.V., Amsterdam, The Netherlands), at 512 Hz. Vertical eye-movements were recorded with electrodes located above and below the left eye, horizontal eye-movements were recorded with electrodes located at the outer canthi of the left and the right eye. Pre-processing of EEG-data was conducted using custom-made MATLAB (R2012b, The MathWorks, Inc.) scripts, supported by EEGLAB functionality (Delorme & Makeig, 2004). EEG-traces were high-pass filtered (>0.1 Hz) to remove slow drifts and re-referenced to the average of two electrodes located on the left and right earlobes (mastoidal reference for 1 participant). Epochs were created by taking data from −1s to 2s around onset of stimulus presentation. Trials containing irregular noise were rejected manually and faulty electrodes were interpolated (spherical interpolation), after which independent components (ICs) were calculated (30 ICs, blind source separation). Components representing eye-blinks, eye-movements, muscle artifacts or other sources of noise were removed. In case these artifacts were still present after IC removal, a second ICA was run over the residual signal. Epochs were grouped based on current and previous trial congruency, creating four trial conditions. Thereafter, the data were CSD-transformed by applying a surface Laplacian filter, thereby increasing spatial specificity (Cohen, 2015). Then, EEG-traces were decomposed into time-frequency representations from 2Hz to 30Hz in 15 linearly spaced steps. The power spectrum of the EEG-signal (as obtained by the fast Fourier transform) was multiplied by the power spectra of complex Morlet wavelets 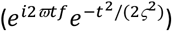 with logarithmically spaced cycle sizes ranging from 3 to 12. The inverse Fourier transform was then used to acquire the complex signal, which was converted to frequency-band specific power by squaring the result of the convolution of the complex and real parts of the signal (*real*[*z*(*t*)]^2^ + *imag*[*z*(*t*)]^2^). The resulting time-frequency data were then resampled to 40Hz, averaged per subject and trial type. Finally, time-frequency traces were transformed to decibels (dB) and normalized to a baseline of −300ms to −100ms before stimulus onset, according to: 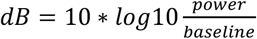 (Cohen & van Gaal, 2013).

### EEG: time-frequency data analyses

As previous literature has shown midfrontal theta-power enhancement following the presentation of conflicting stimuli (Cohen & Ridderinkhof, 2013; Cohen & van Gaal, 2013, 2014; Nigbur et al., 2012; Pastötter et al., 2013) we had a priori hypotheses concerning this electrophysiological signal. Therefore, we preselected a fronto-central spatial region of interest (ROI) to run our analysis on (figure 3A/C). In order to find a time-frequency ROI for subsequent analysis, data from the electrodes within this spatial ROI were averaged over all task-relevance conditions and experimental sessions. Next, current trial conflict was calculated (I-C) for all participants. To extract the time-frequency ROI in which overall conflict effect was present, t-tests were used on this contrast (p<0.01, TF region: −200ms to 1200ms and 2Hz to 30Hz). Cluster-based permutation testing (p<0.01, 1000 permutes) was implemented to correct for multiple comparisons (Maris & Oostenveld, 2007), resulting in a significant time-frequency ROI. Then, time-frequency power was extracted from this ROI for each participant and used as input for rm-ANOVAs with task-relevance, training, current trial congruency and previous trial congruency (2×2×2×2 factorial design). Subsequently, separate rm-ANOVAs for the task-relevant and task-irrelevant conditions were performed on the same ROI data for post-hoc inspection of significant effects containing condition as a factor (2×2×2 factorial design). In the case of null-findings, we performed Bayesian rm-ANOVAs with the same factorial design (JASP Team, 2018).

### EEG: decoding analysis

In addition to this univariate approach, a multivariate backwards decoding model was applied on the time-frequency data. This was done both because of the higher sensitivity of multivariate analyses and to inspect if and to what extent different stimulus features (i.e. location and content) were processed in both conditions. The ADAM-toolbox was used on raw EEG data, that was transformed to time-frequency using default methods of the toolbox but with similar settings (epochs: −200ms to 1200ms, 2Hz-30Hz) (Fahrenfort, Van Driel, van Gaal, & Olivers, 2018). To increase decoding power, time-frequency data of both experimental sessions were merged, thus only separating conditions. Trials were classified according to current trial stimulus features (i.e. location and content) resulting in 4 trial types. Note that this is different from the univariate analyses, where trials were classified according to current and previous trial conflict. As decoding algorithms are known to be time-consuming, data were resampled to 64Hz. Next, a decoding algorithm, using either stimulus location, stimulus content or congruency as stimulus class, was applied on the data according to a tenfold cross-validation scheme. A linear discriminant analysis (LDA) was used to discriminate between stimulus classes (e.g. left versus right speaker location etc.). Classification accuracy was computed as the area under the curve (AUC), a measure derived from Signal Detection Theory (Green & Swets, 1966). AUC scores were tested per time-point and frequency with two-sided t-tests across subjects against a 50% chance-level. These t-test were corrected for multiple comparisons over time, using cluster-based permutation tests (p<0.05, 1000 iterations). This procedure yields time clusters of significant above-chance classifier accuracy, indicative of information processing. Note that this procedure yields results that should be interpreted as fixed effects (Allefeld, Görgen, & Haynes, 2016), but is nonetheless standard in the scientific community.

We used the significant cluster of task-relevant conflict as a mask to extract AUC values. Next, we applied a Bayesian one-sample t-test (one-sided, 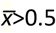, Cauchy scale=0.71) on these AUC values, to confirm the absence of processing of task-irrelevant conflict.

### EEG: Source reconstruction analysis

To visualize the cortical origins of any univariate or multivariate effects we applied a source-reconstruction analysis on our data using the BrainStorm toolbox (Tadel, Baillet, Mosher, Pantazis, & Leahy, 2011). For our univariate data, we performed this analysis on the group-averaged values of individual EEG-channels, from within the theta-band ROI (figure 2). To reconstruct the sources of multivariate effects, individual classifier weights were extracted from the significant clusters and transformed into activation patterns by multiplying them with the covariance in the data (Grootswagers, Wardle, & Carlson, 2017). Subsequently, sources were constructed from these data and averaged across subjects. Because there was no above-chance decoding present when the auditory stimuli were task-irrelevant, we used the cluster of the task-relevant data as a mask to extract AUC-values. However, note that this procedure does not allow for proper statistical testing and therefore should be considered as a visualization tool only.

## Supplementary information and figures

### Descriptive statistics of behavioral results

For the task-relevant condition, error rates (ERs) and reaction times (RTs), averaged over all four sessions, were 2.6% (*s*=2.7%) and 474.2ms (*s*=76.1ms), respectively. Behavioral results for the task-irrelevant condition were different, with mean ERs of 19.2% (*s*=6.6%) a mean RTs of 711.4ms (*s*=151.3ms). The mean ER of the task-irrelevant condition indicates that our staircasing-procedure, 75% correct, was effective.

### Effects of training on relative conflict effect and adaptation

**Figure S1.**
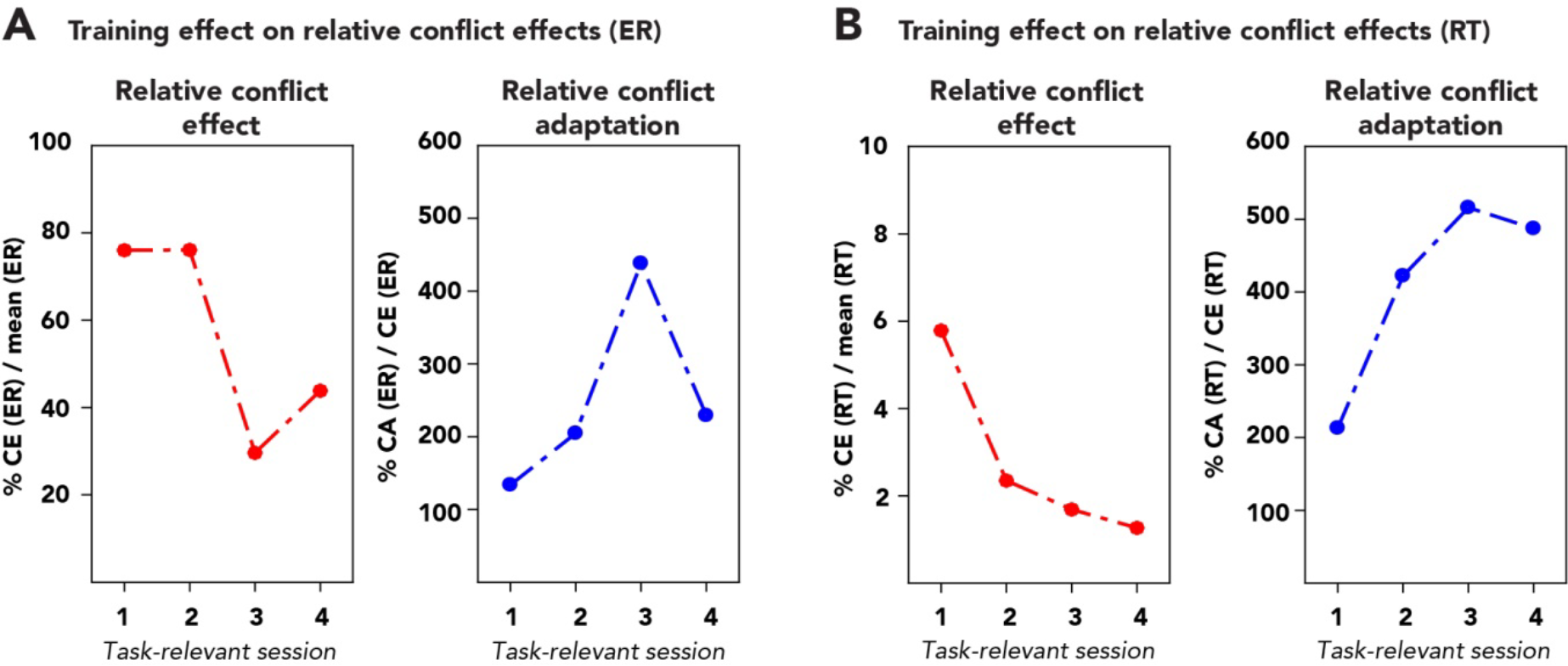
Effects of training on relative conflict effect and adaptation in terms of ER and RT. Relative measures of conflict effect and conflict adaptation provide better insight in the magnitude of effects, albeit unconventional measures. All values are grand averages, and statistics were not applied on them. **(A)** Training effects on relative conflict effect (left) and conflict adaptation (right) for ERs. Whereas relative conflict effect seems to decrease as a result of training, relative conflict adaptation appears to increase. This is suggestive of more efficient conflict resolution. **(B)** Training effects on relative conflict effect (left) and conflict adaptation (right) for RTs. Similar to ERs relative conflict effect decreases as a result of training, whereas relative conflict adaptation appears to increase. Again, this suggests more efficient conflict resolution.

### Decoding performance of before and after training sessions

In our **Results** section we report the outcome of multivariate analyses on EEG-data that was merged over the two experimental sessions. We merged these data to increase classifier performance. Before merging the data, we ran an identical analysis on the session-specific data. Crucially, no differences in congruency classifier performance were found between before and after training data, for neither the task-relevant nor task-irrelevant condition (p>0.05, cluster-corrected; see supplementary figure S2).

**Figure S2.**
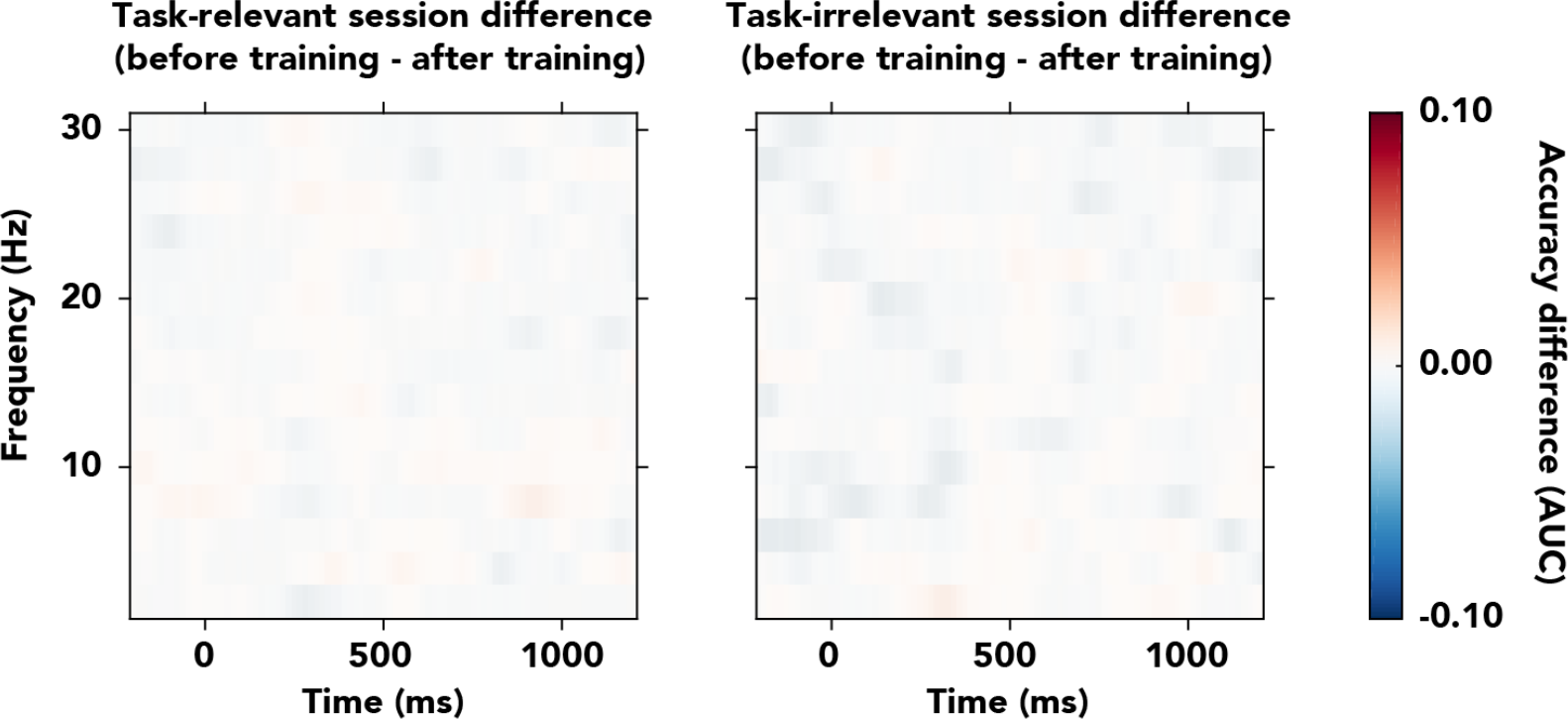
Between session difference (before training – after training) in classifier performance for congruency, for the task-relevant and task-irrelevant condition. The accuracy differences are depicted across the frequency-range (2-30Hz) for the task-relevant condition (left panel) and task-irrelevant condition (right panel). Classifier accuracy differences are thresholded (cluster-base corrected, p<0.05) and non-significant clusters were masked. No significant clusters were observed, indicating that classifier performance was not affected by extensive training in the conflict task, neither for task-relevant (left) nor task-irrelevant (right) auditory stimuli.

### Excluding auditory interference in task-irrelevant condition

Although we report no effects of task-irrelevant auditory conflict, some dimensions of the auditory stimuli could still have interfered with performance on the RDM-task. For example, the spoken word “left” or an auditory stimulus presented from the left side of the participant could lead to conflict with right hand responses in the task-irrelevant condition. To test such potential interference, we performed a 2×2×2 factorial rm-ANOVA on mean RTs of the task-irrelevant condition, with session (before/after training), stimulus content (“left”/”right”), and stimulus location (left/right) as factors. RTs in the task-irrelevant condition were unaffected both by stimulus content (*F*_*1,23*_=0.29, *p*=0.59, 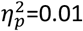, BF_01_=6.42) and location (*F*_*1,23*_=0.41, *p*=0.53, 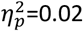, BF_01_=6.33) (see figure S3, data for task-relevant condition plotted as control).

**Figure S3.**
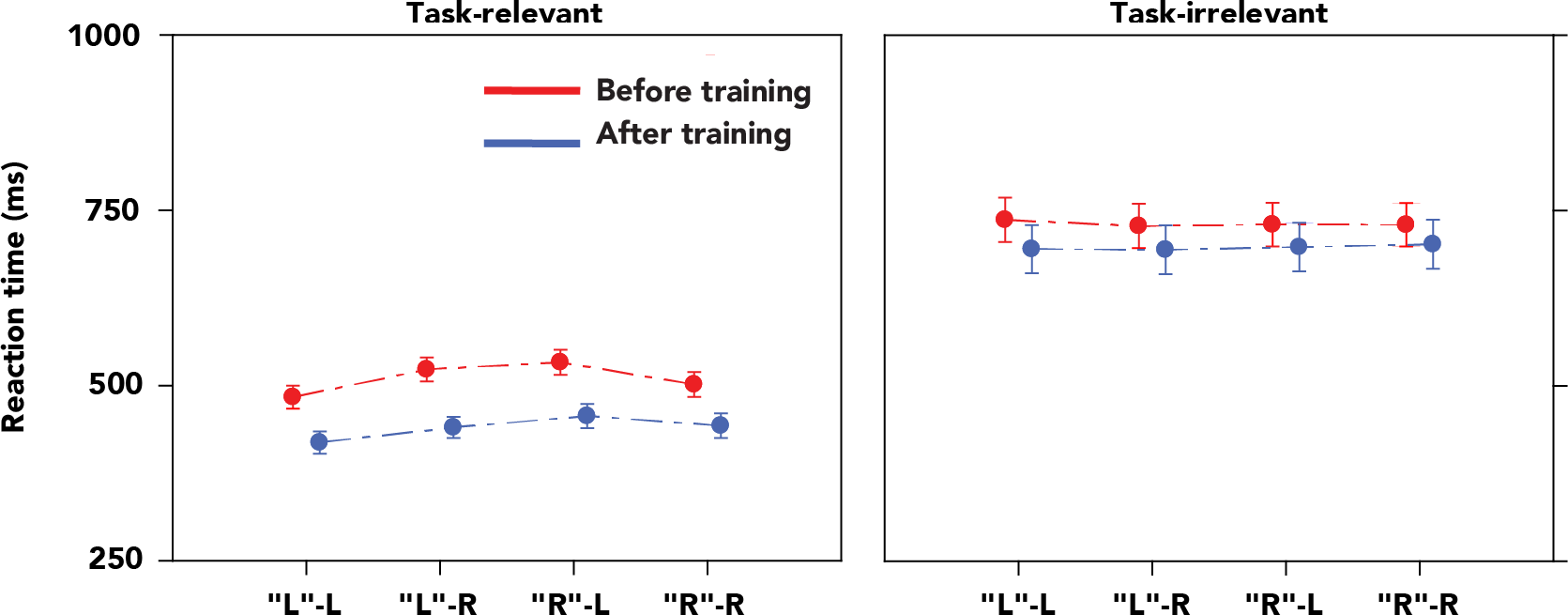
Effect of auditory stimulus on reaction times for both the task-relevant and task-irrelevant condition. When the auditory stimulus was task-relevant, reaction times were affected by the specific auditory stimulus that was presented (left pane). Specifically, reaction times were increased for stimuli of which stimulus content (first letter, L or R denoted in quotation marks) and stimulus location (second letter, L or R) were incongruent. Contrary, reaction times in the task-irrelevant condition were unaffected by the specific auditory stimulus, excluding any possible conflict between stimulus content or location and responses made with exclusively the right hand (right pane).

### Excluding conflict task-related effects on decoding in task irrelevant condition

In our experimental design, we excluded across-trial fluctuations of task-relevance of the auditory stimulus, by applying a blocked design. This ensured that the auditory stimulus in the task-irrelevant condition was task-irrelevant on *every single trial*. However, participants were extensively trained in the auditory conflict task, and as such participants might have been prompt to detect the auditory stimuli in the second session of the task-irrelevant condition, due to their earlier relevance. This in turn might have confounded our decoding results, for which we merged data from both EEG sessions, before and after training in the conflict task. That is, the above chance decoding of stimulus content in the task-irrelevant condition might mainly be caused by some sort of lingering *background* process aiming to detect the previously task-relevant stimulus content of the auditory stimulus. However, stimulus content could already be decoded from data of the first session of the task-irrelevant condition, when participants were not yet exposed to the conflicting task (see figure S4A). Similarly, stimulus location could also be classified above-chance before the auditory stimuli gained any task-relevance (see figure S4B). Thus, the significant decoding of stimulus content and location in the task-irrelevant condition indeed reflects perceptual processing of truly irrelevant stimulus features.

**Figure S4.**
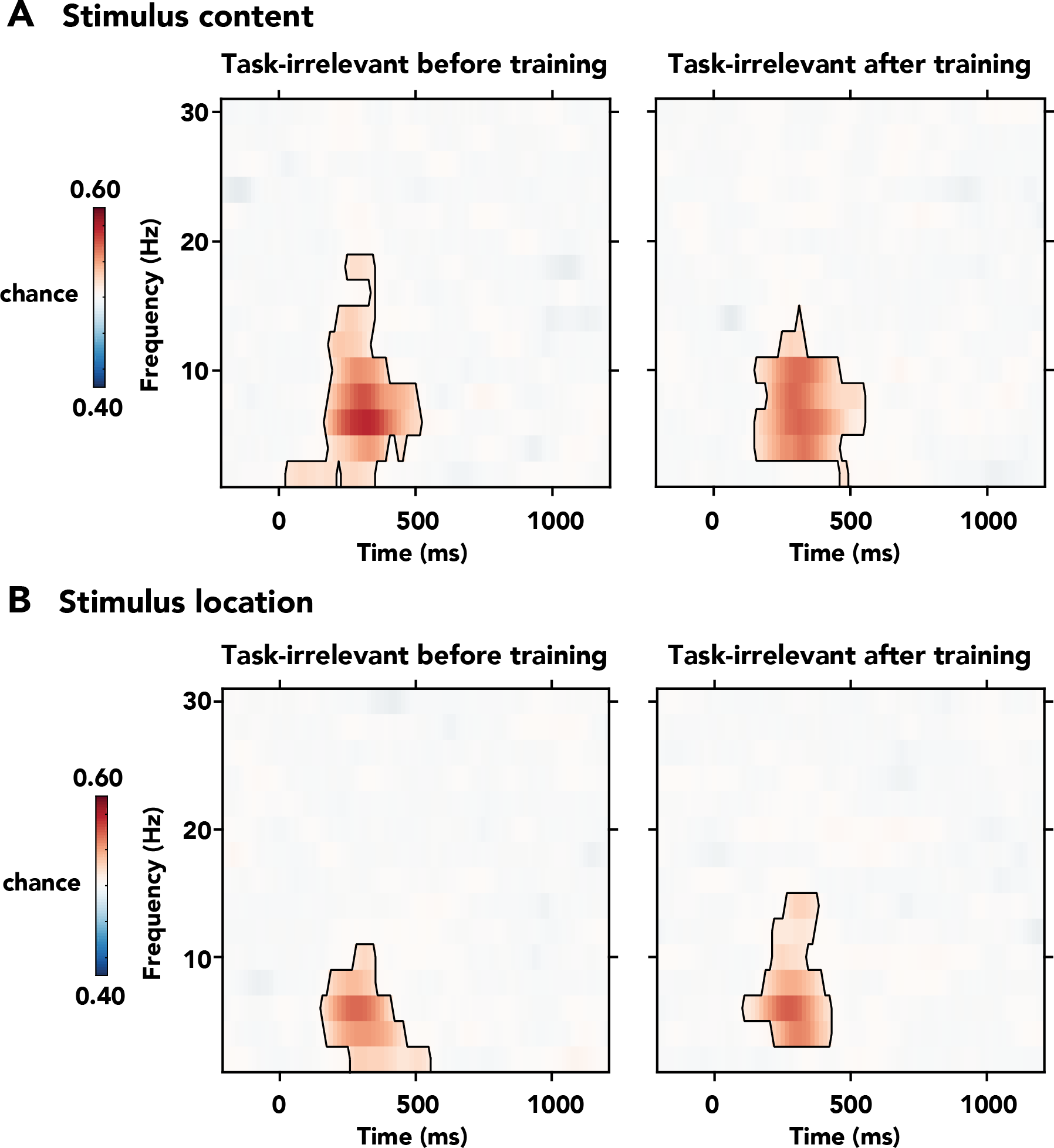
Classifier accuracies for auditory stimulus content, in the task-irrelevant condition before and training in the conflicting task. Thresholded (cluster-base corrected, p<0.05) accuracies are depicted across the frequency-range (2-30Hz) and significant clusters are outlined with a solid black line. Source activations are expressed in arbitrary units (a.u.) and are thresholded at 50% of the within-condition maximum value. **(A/B)** Classifier accuracies of **(A)** stimulus content and **(B)** stimulus location in the task-irrelevant condition before training (left panel) and after training (right panel) in the conflicting task. During the first session of the task-irrelevant condition, participants had not yet performed a single trial of the conflicting task. Therefore, the above-chance classification of stimulus content and location shown here is not confounded by any effects related to the conflicting task and thus shows perceptual processing of truly task-irrelevant stimulus features.

### Exploratory analysis of other neural signatures of conflict

Based on previous observations of conflict-induced modulations of parieto-occipital alpha and centro-parietal beta-band activity (Jiang, Zhang, & van Gaal, 2015; Pastötter et al., 2013), we performed exploratory analysis on these ROIs (figure S5). ANOVAs on the extracted power of these ROIs showed decreased alpha-band power within the pre-defined time-frequency ROI over parieto-occipital regions, following incongruent trials as compared to congruent trials (*F*_*1,23*_=7.90, *p*=0.01). This conflict effect was not modulated by task-relevance (*F*_*1,23*_=0.35, *p*=0.56), although the effect was only significant when the auditory stimulus was task-relevant and not when it was task-irrelevant (alpha_*rel*_: *F*_*1,23*_=5.32, *p*=0.03; alpha_irrel_: *F*_*1,23*_=1.410, *p*=0.25). No main effects of conflict were found in the pre-defined beta-band ROI over centro-parietal regions (all *p*>0.05).

**Figure S5.**
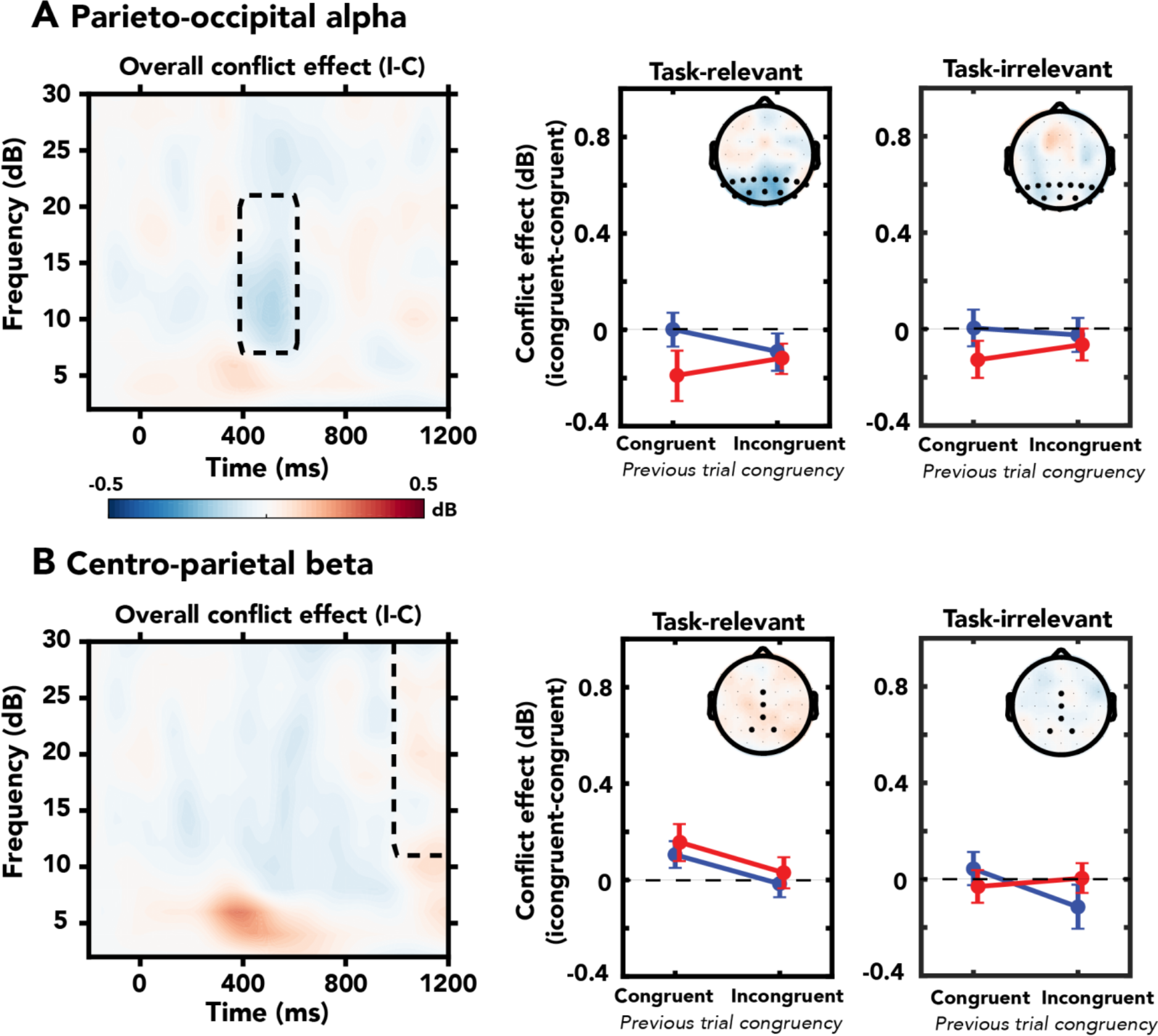
Results of exploratory analysis in the conflict effects in parieto-occipital alpha and centro-parietal beta oscillations. ROIs were based on previous findings of conflict-related effects in these frequency bands. **(A)** A main effect of current trial conflict was found within the pre-selected parieto-occipital ROI (time-range: 400-600ms, frequency-range: 8-20Hz, channels: P1, P2, P3, P4, P5, P6, P7, P8, Pz, PO3, PO4, PO7, PO8, POz, O1, O2, Oz). This conflict effect was not modulated by task-relevance of the auditory stimulus, but nonetheless only observed for task-relevant conflict. **(B)** No conflict-related effects were found within the centro-parietal beta ROI (time-range: 1000-1200ms, frequency-range: 12-30Hz, channels: FCz, Cz, CPz, P1, P2).

## Literature

Allefeld, C., Görgen, K., & Haynes, J. D. (2016). Valid population inference for information-based imaging: From the second-level t-test to prevalence inference. NeuroImage, 141, 378–392. http://doi.org/10.1016/j.neuroimage.2016.07.040

Atas, A., Desender, K., Gevers, W., & Cleeremans, A. (2016). Dissociating perception from action during conscious and unconscious conflict adaptation. Journal of Experimental Psychology: Learning Memory and Cognition. http://doi.org/10.1037/xlm0000206

Botvinick, M. M., Cohen, J. D., & Carter, C. S. (2004). Conflict monitoring and anterior cingulate cortex : an update, 8(12), 8–12. http://doi.org/10.1016/j.tics.2004.10.003

Broadbent, D. E. (1958). Perception and Communication. New York: Oxford University Press.

Buzzell, G. A., Roberts, D. M., Baldwin, C. L., & McDonald, C. G. (2013). An electrophysiological correlate of conflict processing in an auditory spatial Stroop task: The effect of individual differences in navigational style. International Journal of Psychophysiology, 90(2), 265–271. http://doi.org/10.1016/j.ijpsycho.2013.08.008

Canales-Johnson, A., Billig, A., Olivares, F., Gonzalez, A., Garcia, M. del C., Silva, W., … Bekinschtein, T. (2019). Dissociable neural information dynamics of perceptual integration and differentiation during bistable perception. BioRxiv, 133801. http://doi.org/10.1101/133801

Cavanagh, J. F., & Frank, M. J. (2014). Frontal theta as a mechanism for cognitive control. Trends in Cognitive Sciences, 18(8), 414–421. http://doi.org/10.1016/j.tics.2014.04.012

Cherry, E. C. (1953). Some experiments on the recognition of speech, with one and with 2 ears. Journal of the Acoustical Society of America. http://doi.org/10.1121/1.1907229

Cohen, M. X. (2014). A neural microcircuit for cognitive conflict detection and signaling. Trends in Neurosciences, 37(9), 480–490. http://doi.org/10.1016/j.tins.2014.06.004

Cohen, M. X. (2015). Comparison of different spatial transformations applied to EEG data: A case study of error processing. International Journal of Psychophysiology, 97(3), 245–257. http://doi.org/10.1016/j.ijpsycho.2014.09.013

Cohen, M. X., & Cavanagh, J. F. (2011). Single-trial regression elucidates the role of prefrontal theta oscillations in response conflict. Frontiers in Psychology, 2(FEB), 1–12. http://doi.org/10.3389/fpsyg.2011.00030

Cohen, M. X., & Donner, T. H. (2013). Midfrontal conflict-related theta-band power reflects neural oscillations that predict behavior. Journal of Neurophysiology, 110(12), 2752–2763. http://doi.org/10.1152/jn.00479.2013

Cohen, M. X., & Ridderinkhof, K. R. (2013). EEG Source Reconstruction Reveals Frontal-Parietal Dynamics of Spatial Conflict Processing. PLoS ONE, 8(2). http://doi.org/10.1371/journal.pone.0057293

Cohen, M. X., & van Gaal, S. (2013). Dynamic interactions between large-scale brain networks predict behavioral adaptation after perceptual errors. Cerebral Cortex, 23(5), 1061–1072. http://doi.org/10.1093/cercor/bhs069

Cohen, M. X., & van Gaal, S. (2014). Subthreshold muscle twitches dissociate oscillatory neural signatures of conflicts from errors. NeuroImage, 86, 503–513. http://doi.org/10.1016/j.neuroimage.2013.10.033

D’Ostilio, K., & Garraux, G. (2012a). Brain mechanisms underlying automatic and unconscious control of motor action. Frontiers in Human Neuroscience, 6(September), 1–5. http://doi.org/10.3389/fnhum.2012.00265

D’Ostilio, K., & Garraux, G. (2012b). Dissociation between unconscious motor response facilitation and conflict in medial frontal areas. European Journal of Neuroscience, 35(2), 332–340. http://doi.org/10.1111/j.1460-9568.2011.07941.x

Dehaene, S., Changeux, J. P., Naccache, L., Sackur, J., & Sergent, C. (2006). Conscious, preconscious, and subliminal processing: a testable taxonomy. Trends in Cognitive Sciences, 10(5), 204–211. http://doi.org/10.1016/j.tics.2006.03.007

Dehaene, S., & Naccache, L. (2001). Towards a cognitive neuroscience of consciousness: Basic evidence and a workspace framework. Cognition, 79(1-2), 1–37. http://doi.org/10.1016/S0010-0277(00)00123-2

Delorme, A., & Makeig, S. (2004). EEGLAB: An open source toolbox for analysis of single-trial EEG dynamics including independent component analysis. Journal of Neuroscience Methods, 134(1), 9–21. http://doi.org/10.1016/j.jneumeth.2003.10.009

Deutsch, J. A., & Deutsch, D. (1963). Attention: Some theoretical considerations. Psychological Review, 70. http://doi.org/10.1037/h0042712

Egner, T. (2007). Congruency sequence effects. Cognitive, Affective & Behavioral Neuroscience, 7(4), 380–390. http://doi.org/10.3758/CABN.7.4.380

Fahrenfort, J. J., Van Driel, J., van Gaal, S., & Olivers, C. N. L. (2018). From ERPs to MVPA using the Amsterdam Decoding and Modeling toolbox (ADAM). Frontiers in Neuroscience – Brain Imaging Methods, 12(July). http://doi.org/10.3389/fnins.2018.00368

Fahrenfort, J. J., van Leeuwen, J., Olivers, C. N. L., & Hogendoorn, H. (2017). Perceptual integration without conscious access. Proceedings of the National Academy of Sciences, 114(14), 3744–3749. http://doi.org/10.1073/pnas.1617268114

Fei-Fei, L., VanRullen, R., Koch, C., & Perona, P. (2002). Rapid natural scene categorization in the near absence of attention. Proceedings of the National Academy of Sciences, 99(14), 9596–9601. http://doi.org/10.1073/pnas.092277599

Finkbeiner, M., & Palermo, R. (2017). The Role of Spatial Attention in Nonconscious Processing A Comparison of Face and Nonface Stimuli. Psychological Science, 20(1), 42–51.

Frühholz, S., Godde, B., Finke, M., & Herrmann, M. (2011). Spatio-temporal brain dynamics in a combined stimulus-stimulus and stimulus-response conflict task. NeuroImage, 54(1), 622–634. http://doi.org/10.1016/j.neuroimage.2010.07.071

Fuentemilla, L., Marco-Pallarés, J., & Grau, C. (2006). Modulation of spectral power and of phase resetting of EEG contributes differentially to the generation of auditory event-related potentials. NeuroImage, 30(3), 909–916. http://doi.org/10.1016/j.neuroimage.2005.10.036

Giraud, A. L., & Poeppel, D. (2012). Cortical oscillations and speech processing: Emerging computational principles and operations. Nature Neuroscience, 15(4), 511–517. http://doi.org/10.1038/nn.3063

Green, D. M., & Swets, J. a. (1966). Signal detection theory and psychophysics. Society. http://doi.org/10.1901/jeab.1969.12-475

Gronau, N., & Izoutcheev, A. (2017). The necessity of visual attention to scene categorization: Dissociating “task-relevant” and “task-irrelevant” scene distractors. Journal of Experimental Psychology: Human Perception and Performance, 43(5), 954–970. http://doi.org/10.1037/xhp0000365

Grootswagers, T., Wardle, S. G., & Carlson, T. A. (2017). Decoding Dynamic Brain Patterns from Evoked Responses: A Tutorial on Multivariate Pattern Analysis Applied to Time Series Neuroimaging Data. Journal of Cognitive Neuroscience, 29(4), 677–697. http://doi.org/10.1162/jocn_a_01068

Hebart, M. N., & Baker, C. I. (2017). Deconstructing multivariate decoding for the study of brain function. NeuroImage, (August), 1–15. http://doi.org/10.1016/j.neuroimage.2017.08.005

Huber-Huber, C., & Ansorge, U. (2018). Unconscious conflict adaptation without feature-repetitions and response time carry-over. Journal of Experimental Psychology: Human Perception and Performance, 44(2), 169–175. http://doi.org/10.1037/xhp0000450

Itthipuripat, S., Deering, S., & Serences, J. T. (2019). When Conflict Cannot be Avoided: Relative Contributions of Early Selection and Frontal Executive Control in Mitigating Stroop Conflict. Cerebral Cortex, 1–12. http://doi.org/10.1093/cercor/bhz042

Jääskeläinen, I. P., Tapani, K., Lahnakoski, J. M., Manoach, D. S., Sams, M., Ahveninen, J., … Glerean, E. (2016). Neural mechanisms supporting evaluation of others’ errors in real-life like conditions. Scientific Reports, 6(1), 1–10. http://doi.org/10.1038/srep18714

JASP Team. (2018). JASP (Version 0.8.6.0). [Computer Software].

Jiang, J., Correa, C. M., Geerts, J., & van Gaal, S. (2018). The relationship between conflict awareness and behavioral and oscillatory signatures of immediate and delayed cognitive control. NeuroImage, 177(May), 11–19. http://doi.org/10.1016/j.neuroimage.2018.05.007

Jiang, J., Zhang, Q., & van Gaal, S. (2015). Conflict awareness dissociates theta-band neural dynamics of the medial frontal and lateral frontal cortex during trial-by-trial cognitive control. NeuroImage, 116, 102–111. http://doi.org/10.1016/j.neuroimage.2015.04.062

Jiang, J., Zhang, Q., & Van Gaal, S. (2015). EEG neural oscillatory dynamics reveal semantic and response conflict at difference levels of conflict awareness. Scientific Reports, 5(July), 1–11. http://doi.org/10.1038/srep12008

Kleiner, M., Brainard, D. H., & Pelli, D. G. (2007). What’s new in Psychtoobox-3? Perception.

Koch, C., & Tsuchiya, N. (2007). Attention and consciousness: two distinct brain processes. Trends in Cognitive Sciences, 11(1), 16–22. http://doi.org/10.1016/j.tics.2006.10.012

Koelewijn, T., Bronkhorst, A., & Theeuwes, J. (2010). Attention and the multiple stages of multisensory integration: A review of audiovisual studies. Acta Psychologica, 134(3), 372–384. http://doi.org/10.1016/j.actpsy.2010.03.010

Kornblum, S. (1994). The way irrelevant dimensions are processed depends on what they overlap with: The case of Stroop- and Simon-like stimuli. Psychological Research, 56(3), 130–135. http://doi.org/10.1007/BF00419699

Kouider, S., Andrillon, T., Barbosa, L. S., Goupil, L., & Bekinschtein, T. A. (2014). Inducing task-relevant responses to speech in the sleeping brain. Current Biology, 24(18), 2208–2214. http://doi.org/10.1016/j.cub.2014.08.016

Lachter, J., Forster, K. I., & Ruthruff, E. (2004). Forty-five years after broadbent (1958): Still no identification without attention. Psychological Review. http://doi.org/10.1037/0033-295X.111.4.880

Lamme, V. A. F. (2003). Why visual attention and awareness are different. Trends in Cognitive Sciences, 7(1), 12–18. http://doi.org/10.1016/S1364-6613(02)00013-X

Lamme, V. A. F., & Roelfsema, P. R. (2000). The distinct modes of vision offered by feedforward and recurrent processing. Trends in Neurosciences, 1–9. http://doi.org/10.10162236

Lavie, N., Ro, T., & Russell, C. (2003). The role of perceptual load in processing distractor faces. Psychological Science, 14(5), 510–515. http://doi.org/10.1111/1467-9280.03453

Lewald, J., & Getzmann, S. (2011). When and where of auditory spatial processing in cortex: A novel approach using electrotomography. PLoS ONE, 6(9). http://doi.org/10.1371/journal.pone.0025146

Mao, W., & Wang, Y. (2008). The active inhibition for the processing of visual irrelevant conflict information. International Journal of Psychophysiology, 67(1), 47–53. http://doi.org/10.1016/j.ijpsycho.2007.10.003

Maris, E., & Oostenveld, R. (2007). Nonparametric statistical testing of EEG- and MEG-data. Journal of Neuroscience Methods, 164(1), 177–190. http://doi.org/10.1016/j.jneumeth.2007.03.024

Molloy, K., Griffiths, T. D., Chait, M., & Lavie, N. (2015). Inattentional Deafness: Visual Load Leads to Time-Specific Suppression of Auditory Evoked Responses. Journal of Neuroscience, 35(49), 16046–16054. http://doi.org/10.1523/JNEUROSCI.2931-15.2015

Moray, N. (1959). Attention in Dichotic Listening: Affective Cues and the Influence of Instructions. Quarterly Journal of Experimental Psychology, 11(1). http://doi.org/10.1080/17470215908416289

Mudrik, L., Faivre, N., & Koch, C. (2014). Information integration without awareness. Trends in Cognitive Sciences, 18(9), 488–496. http://doi.org/10.1016/j.tics.2014.04.009

Nigbur, R., Cohen, M. X., Ridderinkhof, K. R., & Stürmer, B. (2012). Theta Dynamics Reveal Domain-specific Control over Stimulus and Response Conflict. Journal of Cognitive Neuroscience, 24(5), 1264–1274. http://doi.org/10.1162/jocn_a_00128

Padrão, G., Rodriguez-Herreros, B., Pérez Zapata, L., & Rodriguez-Fornells, A. (2015). Exogenous capture of medial-frontal oscillatory mechanisms by unattended conflicting information. Neuropsychologia, 75, 458–468. http://doi.org/10.1016/j.neuropsychologia.2015.07.004

Pastötter, B., Dreisbach, G., & Bäuml, K.-H. T. (2013). Dynamic Adjustments of Cognitive Control: Oscillatory Correlates of the Conflict Adaptation Effect. Journal of Cognitive Neuroscience, 25(12), 2167–2178. http://doi.org/10.1162/jocn_a_00474

Peelen, M. V., Fei-Fei, L., & Kastner, S. (2009). Neural mechanisms of rapid natural scene categorization in human visual cortex. Nature, 460(7251), 94–97. http://doi.org/10.1038/nature08103

Rahnev, D. A., Huang, E., & Lau, H. C. (2012). Subliminal stimuli in the near absence of attention influence top-down cognitive control. Attention, Perception, and Psychophysics, 74(3), 521–532. http://doi.org/10.3758/s13414-011-0246-z

Ridderinkhof, K. R., Ullsperger, M., Crone, E. A., & Nieuwenhuis, S. (2004). The role of the medial frontal cortex in cognitive control. Science, 306(5695), 443–447. http://doi.org/10.1126/science.1100301

Röer, J. P., Körner, U., Buchner, A., & Bell, R. (2017). Semantic priming by irrelevant speech. Psychonomic Bulletin and Review, 24(4), 1205–1210. http://doi.org/10.3758/s13423-016-1186-3

Rousselet, G. A., Thorpe, S. J., & Fabre-Thorpe, M. (2004). How parallel is visual processing in the ventral pathway? Trends in Cognitive Sciences, 8(8), 363–370. http://doi.org/10.1016/j.tics.2004.06.003

Salminen, N. H., Takanen, M., Santala, O., Lamminsalo, J., Altoè, A., & Pulkki, V. (2015). Integrated processing of spatial cues in human auditory cortex. Hearing Research, 327, 143–152. http://doi.org/10.1016/j.heares.2015.06.006

Sand, A., & Wiens, S. (2011). Processing of unattended, simple negative pictures resists perceptual load. NeuroReport, 22(7), 348–352. http://doi.org/10.1097/WNR.0b013e3283463cb1

Schnuerch, R., Kreitz, C., Gibbons, H., & Memmert, D. (2016). Not quite so blind: Semantic processing despite inattentional blindness. Journal of Experimental Psychology: Human Perception and Performance, 42(4), 459–463. http://doi.org/10.1037/xhp0000205

Sergent, C., Baillet, S., & Dehaene, S. (2005). Timing of the brain events underlying access to consciousness during the attentional blink. Nature Neuroscience, 8(10), 1391–1400. http://doi.org/10.1038/nn1549

Simons, D. J., & Chabris, C. F. (1999). Gorillas in Our Midst: Sustained Inattentional Blindness for Dynamic Events. Perception, 28(9), 1059–1074. http://doi.org/10.1068/p281059

Soutschek, A., Stelzel, C., Paschke, L., Walter, H., & Schubert, T. (2015). Dissociable Effects of Motivation and Expectancy on Conflict Processing: An fMRI Study. J Cogn Neurosci, 27(2), 409–423. http://doi.org/10.1162/jocn_a_00712

Stefanics, G., Csukly, G., Komlósi, S., Czobor, P., & Czigler, I. (2012). Processing of unattended facial emotions: A visual mismatch negativity study. NeuroImage, 59(3), 3042–3049. http://doi.org/10.1016/j.neuroimage.2011.10.041

Tadel, F., Baillet, S., Mosher, J. C., Pantazis, D., & Leahy, R. M. (2011). Brainstorm: A user-friendly application for MEG/EEG analysis. Computational Intelligence and Neuroscience, 2011. http://doi.org/10.1155/2011/879716

Treisman, A. M., & Gelade, G. (1980). A feature-integration theory of attention. Cognitive Psychology, 12(1), 97–136. http://doi.org/10.1016/0010-0285(80)90005-5

Tusche, A., Kahnt, T., Wisniewski, D., & Haynes, J. D. (2013). Automatic processing of political preferences in the human brain. NeuroImage, 72, 174–182. http://doi.org/10.1016/j.neuroimage.2013.01.020

Ullsperger, M., Danielmeier, C., & Jocham, G. (2014). Neurophysiology of Performance Monitoring and Adaptive Behavior. Physiological Reviews, 94(1), 35–79. http://doi.org/10.1152/physrev.00041.2012

Usami, K., Matsumoto, R., Kunieda, T., Shimotake, A., Matsuhashi, M., Miyamoto, S., … Ikeda, A. (2013). Pre-SMA actively engages in conflict processing in human: A combined study of epicortical ERPs and direct cortical stimulation. Neuropsychologia, 51(5), 1011–1017. http://doi.org/10.1016/j.neuropsychologia.2013.02.002

van Gaal, S., de Lange, F. P., & Cohen, M. X. (2012). The role of consciousness in cognitive control and decision making. Frontiers in Human Neuroscience, 6(May), 1–15. http://doi.org/10.3389/fnhum.2012.00121

van Gaal, S., Lamme, V. A. F., Fahrenfort, J. J., & Ridderinkhof, K. R. (2010). Dissociable Brain Mechanisms Underlying the Conscious and Unconscious Control of Behavior. J.Cogn Neurosci., (0898–929X (Linking)), 91–105. http://doi.org/10.1162/jocn.2010.21431

van Gaal, S., Lamme, V. A. F., & Ridderinkhof, K. R. (2010). Unconsciously triggered conflict adaptation. PLoS ONE. http://doi.org/10.1371/journal.pone.0011508

van Gaal, S., Naccache, L., Meuwese, J. D. I., van Loon, A. M., Leighton, A. H., Cohen, L., & Dehaene, S. (2014). Can the meaning of multiple words be integrated unconsciously? Philosophical Transactions of the Royal Society B: Biological Sciences, 369(1641), 20130212–20130212. http://doi.org/10.1098/rstb.2013.0212

van Gaal, S., Ridderinkhof, K. R., Fahrenfort, J. J., Scholte, H. S., & Lamme, V. A. F. (2008a). Frontal Cortex Mediates Unconsciously Triggered Inhibitory Control. Journal of Neuroscience, 28(32), 8053–8062. http://doi.org/10.1523/JNEUROSCI.1278-08.2008

van Gaal, S., Ridderinkhof, K. R., Fahrenfort, J. J., Scholte, H. S., & Lamme, V. A. F. (2008b). Frontal Cortex Mediates Unconsciously Triggered Inhibitory Control. Journal of Neuroscience, 28(32), 8053–8062. http://doi.org/10.1523/JNEUROSCI.1278-08.2008

van Gaal, S., Ridderinkhof, K. R., van den Wildenberg, W. P. M., & Lamme, V. A. F. (2009). Dissociating Consciousness From Inhibitory Control: Evidence for Unconsciously Triggered Response Inhibition in the Stop-Signal Task. Journal of Experimental Psychology: Human Perception and Performance, 35(4), 1129–1139. http://doi.org/10.1037/a0013551

Van Schie, H. T., Mars, R. B., Coles, M. G. H., & Bekkering, H. (2004). Modulation of activity in medial frontal and motor cortices during error observation. Nature Neuroscience, 7(5), 549–554. http://doi.org/10.1038/nn1239

van Veen, V., Cohen, J. D., Botvinick, M. M., Stenger, V. A., & Carter, C. S. (2001). Anterior Cingulate Cortex, Conflict Monitoring, and Levels of Processing. NeuroImage, 14(6), 1302–1308. http://doi.org/10.1006/nimg.2001.0923

VanRullen, R. (2007). The power of the feed-forward sweep. Advances in Cognitive Psychology, 3(1–2), 167–176. http://doi.org/10.2478/v10053-008-0022-3

Wang, K., Li, Q., Zheng, Y., Wang, H., & Liu, X. (2014). Temporal and spectral profiles of stimulus-stimulus and stimulus-response conflict processing. NeuroImage, 89, 280–288. http://doi.org/10.1016/j.neuroimage.2013.11.045

Weisz, N., Müller, N., Jatzev, S., & Bertrand, O. (2014). Oscillatory alpha modulations in right auditory regions reflect the validity of acoustic cues in an auditory spatial attention task. Cerebral Cortex, 24(10), 2579–2590. http://doi.org/10.1093/cercor/bht113

Wolfe, J. M., & Horowitz, T. S. (2004). What attributes guide the deployment of visual attention and how do they do it? Nature Reviews Neuroscience. http://doi.org/10.1038/nrn1411

Zäske, R., Perlich, M. C., & Schweinberger, S. R. (2016). To hear or not to hear: Voice processing under visual load. Attention, Perception, and Psychophysics, 78(5), 1488–1495. http://doi.org/10.3758/s13414-016-1119-2

Zhao, J., Liang, W. K., Juan, C. H., Wang, L., Wang, S., & Zhu, Z. (2015). Dissociated stimulus and response conflict effect in the Stroop task: Evidence from evoked brain potentials and brain oscillations. Biological Psychology, 104, 130–138. http://doi.org/10.1016/j.biopsycho.2014.12.001

Zimmer, U., Itthipanyanan, S., Grent-’T-Jong, T., & Woldorff, M. G. (2010). The electrophysiological time course of the interaction of stimulus conflict and the multisensory spread of attention. European Journal of Neuroscience, 31(10), 1744–1754. http://doi.org/10.1111/j.1460-9568.2010.07229.x

